# SCA44- and SCAR13-associated *GRM1* mutations affect metabotropic glutamate receptor 1 function through distinct mechanisms

**DOI:** 10.1101/2023.07.05.545810

**Authors:** Yuyang Wang, Ashwin Muraleetharan, Karen J Gregory, Shane D Hellyer

## Abstract

Metabotropic glutamate receptor 1 (mGlu_1_) is a promising therapeutic target for neurodegenerative CNS disorders including spinocerebellar ataxias (SCAs). Clinical reports have identified naturally-occurring mGlu_1_ mutations in rare SCA subtypes and clinical symptoms of mGlu_1_ mutations have been described. However, how mutations alter mGlu_1_ function remains unknown. We explored SCA-associated mutation effects on mGlu_1_ cell surface expression and canonical signal transduction. Orthosteric agonists and positive allosteric modulators (PAMs) and negative allosteric modulators (NAMs) were assessed at two functional endpoints (iCa^2+^ mobilisation and IP_1_ accumulation). mGlu_1_ mutants exhibited differential impacts on receptor expression, with a truncating C-terminus mutation significantly reducing mGlu_1_ expression. Mutations differentially influenced orthosteric ligand affinity, efficacy, and functional cooperativity between allosteric and orthosteric ligands. Loss-of-function mutations L454F and N885del reduced orthosteric affinity and efficacy, respectively. Gain-of-function Y792C mutant mGlu_1_ displayed enhanced constitutive activity in IP_1_ assays, which manifested as reduced orthosteric agonist activity. mGlu_1_ PAMs restored glutamate potency in iCa^2+^ mobilisation for loss-of-function mutations, and mGlu_1_ NAMs displayed enhanced inverse agonist activity at Y792C relative to wild-type mGlu_1_. Collectively, these data highlight distinct mechanisms by which mGlu_1_ mutations affect receptor function and show allosteric modulators may present a means to restore aberrant mGlu_1_ function in rare SCA subtypes.

## 1. Introduction

Spinocerebellar ataxia (SCA) describes a heterogeneous range of >40 hereditary neurodegenerative disorders where progressive loss of coordination and balance are the primary symptoms ^1,2^. Progressive cerebellar atrophy is caused by damage to Purkinje neurons, which act as the primary neuronal output of cerebellar cortex ^3,4^. Other tissues such as the spinal cord can also be affected, but the exact pathophysiology and symptomology varies between SCA subtypes ^1^. There are currently no disease modifying treatments for SCA, with clinical failures likely stemming from the genetic diversity of SCAs. The most common SCA subtypes are caused by repeat expansion mutations, resulting in the translation of long glutamine stretches disrupting protein function ^5^. However, some SCA subtypes are caused by conventional (i.e. missense, insertion and deletion) mutations in proteins with important structural or signalling roles in neurons ^1,2^. Given the broad range of causative targets and mutations, the specific dysfunctions resulting from SCA-associated mutations need characterisation to determine potential novel therapies for rare SCA subtypes.

One such SCA-associated signalling protein is mGlu_1,_ one of eight G protein coupled receptors (GPCRs) that respond to the primary excitatory neurotransmitter, glutamate, in the mammalian central nervous system ^6^. An obligate dimer, mGlu_1_ comprises a large extracellular, N-terminal orthosteric ligand binding domain, linked through a cysteine rich domain to the GPCR-characteristic 7 transmembrane (7TM) domain and an extended intracellular C-terminus. mGlu_1_ predominantly couples to G_q/11_ proteins, activating phospholipase C (PLC) to generate inositol 1,4,5-trisphosphate (IP_3_), and subsequently release intracellular Ca^2+^ (iCa^2+^) ^7^. mGlu_1_ is highly expressed in cerebellar Purkinje neurons, where mGlu_1_-iCa^2+^ signalling has critical roles in motor learning and cerebellar development ^8–11^. Abnormal mGlu_1_ function is associated with SCA pathology. Some rare SCA subtypes, including SCA44 (OMIM#617691) and autosomal recessive SCA13 (SCAR13; OMIM#614831), arise from mutations in the *GRM1* gene, which encodes mGlu_1_ ^12–19^. Additionally, models of both repeat-expansion and conventional mutation SCAs have altered mGlu_1_ expression and signalling, indicating mGlu_1_ dysfunction may be a common feature of distinct SCA subtypes ^20–25^.

Encouragingly, altered mGlu_1_ function in both cellular and animal models can be rescued by small molecule allosteric modulators targeting mGlu_1_ ^21,24^. Allosteric modulators potentiate or inhibit agonist activity through a property known as cooperativity, and are categorised as positive allosteric modulator (PAMs) or negative allosteric modulator (NAMs), respectively ^26^. Allosteric modulators offer greater selectivity among mGlu subtypes and also impose a ‘ceiling’ on the magnitude of the allosteric effect, offering greater efficacy and safety compared to traditional therapeutic approaches. Re-purposing existing drugs may also be possible, as the FDA approved drug nitazoxanide was suggested to have mGlu_1_ NAM activity and can modulate SCA-associated mutant mGlu_1_ receptor function in non-canonical measures of function ^14,27^. Therefore, allosteric modulation of mGlu_1_ may represent a viable therapeutic approach in treating genetically heterogenous SCA subtypes.

Minimal functional characterisation of SCA-associated mutant mGlu_1_ receptors has been undertaken, with no prior studies investigating the effects of mutations on canonical IP_3_-iCa^2+^ signalling. Therefore, the exact effects of mutations on mGlu_1_ function remains unknown. Additionally, the ability of allosteric modulators to modulate mutant mGlu_1_ function remains understudied. Here, we characterised the effects of SCA-associated mGlu_1_ mutants on receptor expression and function in inducible HEK293A cell lines using two canonical measures of receptor function: iCa^2+^ mobilisation and IP_1_ accumulation. Ligand affinity, cooperativity, potency and efficacy estimates were derived for structurally diverse orthosteric agonists, PAMs and NAMs. We observed differential effects on mGlu_1_ expression and signalling across the two functional measures, with loss- and gain-of-function occurring through changes to both agonist affinity and efficacy. The putative gain-of-function mutant mGlu_1_-Y792C resulted in reduced agonist potency and efficacy in IP_1_ accumulation, which is linked to enhanced constitutive activity. These data enhance our understanding of SCA-associated mutant mGlu_1_ function, and provides further evidence surrounding the therapeutic potential of mGlu_1_ allosteric modulators in restoring aberrant mGlu_1_ function in rare SCAs.

## 2. Materials and Methods

### 2.1. Materials

Dulbecco’s modified Eagle’s medium (DMEM), Fluo-8-AM, Flp-In TRex HEK293 cell line, tetracycline free FBS (fetal bovine serum) (< 10 ng/ml tetracycline) and blasticidin S HCl were purchased from Invitrogen (Carlsbad, CA, USA). Cisbio IP-One HTRF assay kit was purchased from Genesearch (Arundel, QLD, Australia). 3,4-Dihydro-2H-pyrano[2,3-b]quinolin-7-yl)-(cis-4-methoxycyclohexyl)-methanone (JNJ 16259685) was purchased from HelloBio (Princeton, NJ, USA). N-[4-Chloro-2-[(1,3-dihydro-1,3-dioxo-2H-isoindol-2-yl)methyl]phenyl]-2-hydroxybenzamide (CPPHA) was purchased from Focus Bioscience (QLD, Australia). (9H-xanthen-9-ylcarbonyl)-carbamic acid butyl ester (Ro 67-4853) was purchased from Abcam (Cambridge, UK). 3-(3,5-dioxo-1,2,4-oxadiazolidin-2-yl)-L-alanine (quisqualate) was purchased from Merck (Darmstadt, Germany). AF647-conjugated 9E10 antibody was prepared in-house as described previously (Cook et al., 2015). Unless otherwise stated, all other reagents were purchased from Sigma-Aldrich (St. Louis, MO, US) and were of analytical grade.

### 2.2. Cell culture

Wild-type human mGlu_1_ construct was created as described previously (Muraleetharan et al., 2023). A c-Myc epitope was introduced between residues L384 and L385 using overlap PCR (primers listed in Table S1). Human cMyc-mGlu_1_ cDNA sequence was subsequently transferred from pcDNA3.1(+) into the destination vector pcDNA5/FRT/TO (Invitrogen) using NEBuilder HiFi assembly (New England Biolabs, Notting Hill, Australia). SCA-associated mutations were introduced into human cMyc-mGlu_1_ using Quikchange site-directed mutagenesis (Agilent Technologies, Santa Clara, CA). Mutagenesis primers are shown in Table S1. Successful mutagenesis was confirmed by Sanger sequencing (Australian Genome Research Facility, Melbourne, Australia). To generate stable Flp-In HEK293-TREx c-Myc-mGlu_1_ cells, Flp-In HEK293-TREx cells were transfected with pcDNA5/FRT/TO-cMyc-mGlu_1,_ Lipofectamine 2000 and the pOG44 Flp-recombinase expression vector (Invitrogen) and selected in 200 µg/ml hygromycin B and 5 µg/ml blasticidin. Cells were subsequently maintained in DMEM with 5% tetracycline-free FBS, 200μg/mL hygromycin B and 5 μg/ml blasticidin. Cells were routinely checked for mycoplasma contamination, and were not used past passage 30.

### 2.3. iCa^2+^ mobilisation assays

The day prior to assays, Flp-In HEK293-TREx cells or non-transfected HEK293A cells were seeded onto poly-D-lysine coated, clear or black walled and clear-bottom 96-well plates in assay medium (glutamine-free DMEM supplemented with 2.5% dialysed FBS and 16 mM HEPES) at a density of 80,000 cells/well. For receptor titration assays, Flp-In HEK293-TREx cells expressing mutant and wild type mGlu_1_ were incubated overnight in the presence of 0, 3ng/ml, 10ng/ml, 30ng/ml, 100ng/ml, 300ng/ml, and 1µg/ml of tetracycline to induce different levels of receptor expression. For agonism and modulation studies, receptor expression was induced by overnight treatment with 30ng/ml tetracycline. All iCa^2+^ mobilisation assays were performed in calcium assay buffer (HBSS: 1.2 mM CaCl_2_, KCl 5.33 mM, KH_2_PO_4_ 0.44 mM, NaCl 137.93 mM, Na_2_HPO_4_ 0.34 mM, D-glucose 5.56 mM; supplemented with 20mM HEPES and 4mM probenecid, pH 7.4). Cells were washed with calcium assay buffer and incubated with the cell permeable Ca^2+^ sensitive Fluo-8-AM dye diluted in calcium assay buffer for 45 min at 37°C in a CO_2_ free incubator. Cells were washed with calcium assay buffer again and a FlexStation I or III (Molecular Devices) or Functional Drug Screening System (FDSS; Hamamatsu) was used to assay receptor-mediated iCa^2+^ mobilisation as described previously (Gregory et al., 2012). A “single add” protocol was used to assay receptor titration. For modulation studies, a “double add” paradigm was performed where allosteric ligands were added 1 min prior to the addition of orthosteric ligands. A 20 sec baseline was recorded prior to any addition and the peak fluorescence was defined as difference between maximum response and baseline over 60 sec. Relative fluorescence units were normalised to the response to ionomycin (1µM).

### 2.4. IP_1_ accumulation assays

Flp-In HEK293-TREx cells were seeded onto poly-D-lysine coated, clear-bottom 96-well plates in assay medium at a density of 40,000 cells/well. After incubation overnight with 30ng/ml tetracycline, cells were washed with phosphate buffered saline (PBS) and incubated for 1 hour at 37°C in a CO_2_ free incubator with IP_1_ assay buffer (HBSS, supplemented with 20mM HEPES, 30mM LiCl_2_ at pH 7.4; with 10 U/ml glutamic pyruvic transaminase (GPT) and 10 mM sodium pyruvate to eliminate ambient glutamate). Orthosteric and/or allosteric ligands were added, cells incubated for 1 hour at 37 °C (no CO_2_) and were subsequently lysed with IP_1_ lysis buffer. IP_1_ levels were determined using the Cisbio HTRF® IP-one assay kit per the manufacturer’s instructions and measured using an Envision plate reader (PerkinElmer). Data are expressed as fold of basal IP_1_ levels.

### 2.5. Flow cytometry analysis for receptor expression

Flp-In HEK293-TREx cells were seeded onto clear-bottom 24 well plates in assay medium at a density of 100,000 cells/well and incubated with 0, 30ng/ml or 1µg/ml tetracycline overnight. The following day, cells were harvested with FACs buffer (phosphate-buffered saline supplemented with 0.1% bovine serum albumin (BSA), 2 mM EDTA, and 0.05% NaN_3_) and transferred to 1.5ml tubes. Cells were centrifuged for 4 min at 400 x*g*, 4°C, and resuspended again in either FACs buffer or FACs permeabilisation buffer (FACs buffer as above supplemented with 0.5% Tween). Cells were incubated on ice for 30 min and harvested by centrifuge for 4 min at 400 x*g*, 4°C, prior to resuspend in FACs buffer containing 1 μg/mL AF647-conjugated anti-cMyc 9E10 antibody. Cells were incubated for 30 min at 4°C and resuspended in FACs buffer containing 1 μg/mL propidium iodide. Receptor cell surface expression was quantified using a FACs Canto (Becton Dickinson, Franklin Lakes, New Jersey) via detection of fluorescent antibody bound to live cells. All data were normalised to negative control cells expressing no mGlu_1_ receptors.

### 2.6. Data analysis

All data analysis was performed using GraphPad Prism 9 (GraphPad Software, California, USA).

Agonist-concentration response curves at different receptor expression level induced by different tetracycline concentration were fitted to a model of receptor depletion:

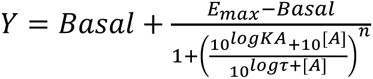

Where [A] denotes the log molar concentration of agonist. E_max_ is the maximum possible system response and basal is the response in the absence of agonist. n represents the transducer slope and was constrained to 1. τ is the transducer constant, an index of agonist efficacy. K_A_ is the agonist equilibrium dissociation constant. To facilitate curve fitting, E_max_ values were constrained to the largest E_max_ value for any mutant/agonist combination (64.22; glutamate at Y792C)

Agonist-concentration response curves at fixed receptor expression level induced by 30ng/ml tetracycline were fitted to a variable four-parameter logistic equation.

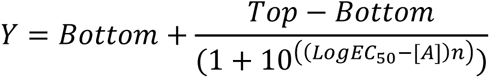

Where bottom and top are the minimum and maximum observed responses, respectively. [A] is the log molar concentration of agonist and n is the Hill coefficient. EC_50_ denotes the agonist concentration required to produce 50% of the maximum response between the top and bottom asymptotes.

Allosteric modulator concentration-response curves in the absence and presence of EC_20_/EC_80_ concentrations of mGlu_1_ orthosteric ligands were fitted to an operational model of allosterism:

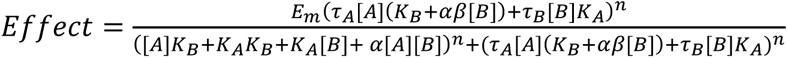

where *E_m_* is the system maximal response, τ_A_ and τ_B_ are the transduction coefficients of orthosteric (A) and allosteric (B) ligands, respectively. α represents affinity cooperativity and β is a scaling factor representing the effect an allosteric modulator has on orthosteric agonist efficacy. For NAMs that fully inhibited responses, β was constrained to −100. K_A_ and K_B_ are the functional affinities of orthosteric and allosteric ligands, respectively. K_A_ was constrained to values derived from receptor titration analysis for each mutant/agonist combination. *n* is the slope factor of the transducer function.

Affinity, cooperativity, potency and efficacy parameters were derived and represented as logarithmic mean ± S.E.M. Statistical analysis of expression level, affinity, potency, and efficacy parameters was performed using a one-way analysis of variance (ANOVA) with Sidak’s post-test to compare mutant receptor responses to WT mGlu_1_, and to compare parameters between assays/ligands for each mutant. All post-test comparisons were chosen before data were observed. Statistical analyses of allosteric modulation data were performed as indicated using an extra sum-of-squares F test to determine the preferred model (logβ unconstrained or neutral (i.e. =0)) for each data set.

## 3. Results

The SCA44 and SCAR13-associated mGlu_1_ mutations p.Tyr262Cys, p.Tyr792Cys, p.Gly1056Argfs*49, p.Asn885del and p.Leu454Phe were chosen for functional analysis in the current study and are referred to throughout as Y262C, Y792C, G1056Rfs*49, N885del and L454F, respectively (Figure 1). The resulting clinical phenotypes and previous experimental data pertaining to changes in mGlu_1_ receptor function are summarised in Table 1. Y262C, L454F are single point mutations in the mGlu_1_ extracellular domain Venus Flytrap domain (VFD), and Y792C is a single point mutation in transmembrane helix 6 of the 7-transmembrane domain (7TM). N885del results in the deletion of an asparagine in the proximal C-terminal tail. It should be noted the N885del mutation co-occurs in patients with an intron 8 splicing mutation ^12^. However, in those patients the functional consequence of the Asn deletion is unclear due to the dominant effect of the splicing. Given we used the alpha splice variant of mGlu_1_ in the current study, and the difficulty of generating complex splice variants with standard DNA manipulation techniques, only the deletion was tested for effects on receptor function. G1056Rfs*49 results in a point mutation of glycine to arginine, and additionally causes a frameshift in the mGlu_1_ C-terminus, resulting in a novel 40 amino acid sequence followed by a premature stop-codon and subsequent truncation of the C-terminal tail. These changes combine to result in a loss of a critical 140 amino acid C-terminal sequence when compared to wild-type (Figure 1).

**Figure 1.**
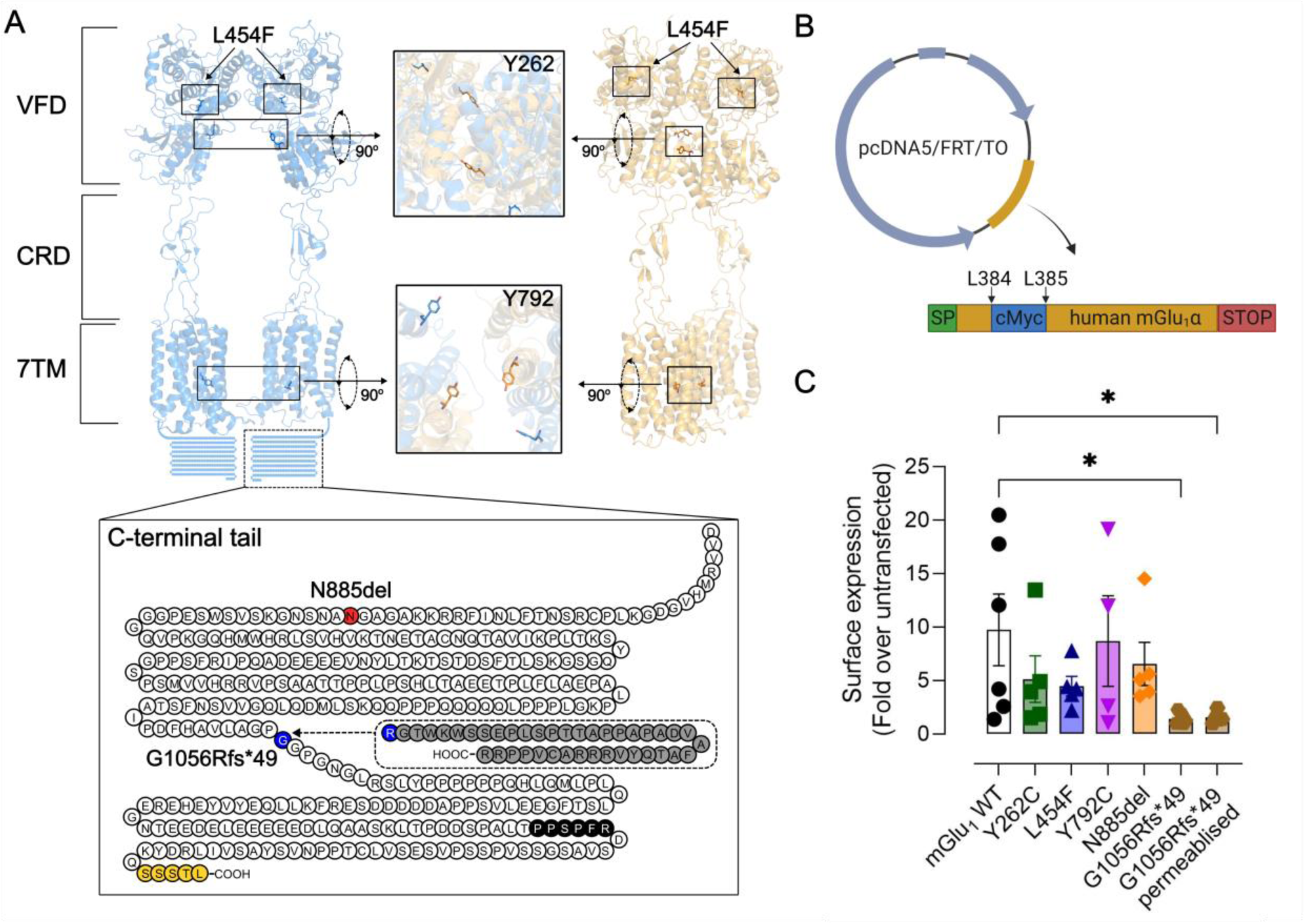
Structural location of mGlu_1_ mutations found in SCA patients and effects of mutations on receptor expression. (A) Cryo-electron microscopy structures of inactive (blue; PDB ID: 7DGD) and active (orange; PDB ID: 7DGE) mGlu_1_ showing the locations of L454F, Y262C and Y792C mutations. Insets for Y262C and Y792C show inward movement of each residue in the active structure relative to the inactive structure. Note the C-terminal tail is not resolved in either structure, and is represented here as a snakeplot. N885del is highlighted in red and G1056R is in blue. The frameshift caused by G1056Rfs*49 is predicted to result in loss of the WT C-tail (in white) and expression of a novel, truncated C-terminus (in grey). As a result, Homer (black) and Tamalin (yellow) binding motifs are predicted to be lost. (B) Map of receptor constructs used in the current study. Human mGlu_1_α isoform in the pcDNA5/FRT/TO vector was modified to incorporate a cMyc epitope between L384 and L385, and mutations were introduced into this tagged receptor. The vector image was created using Biorender.com (C) G1056Rfs*49 significantly reduces cell surface expression relative to WT. Surface expression of mGlu_1_ receptor constructs was detected using FACS with an anti-cMyc antibody following overnight induction of receptor expression in FlpIn TRex HEK293 cells with 30ng/ml tetracycline. Expression was first normalised to non-transfected Flp-In TREx HEK293 cells and then expressed as fold-over 0ng/ml tetracycline for each receptor variant. Data are mean ± S.E.M. pooled from 4-6 or more independent experiments. Significance was determined using an ordinary one-way ANOVA test with Dunnett’s post-hoc test.

**Table 1.** Genetic variants, clinical characteristics and functional effects of spinocerebellar ataxia-associated *GRM1* mutations.

| Nucleotide change | Protein change | SCA subtype | Phenotype | Predicted functional effect | Experimental functional data | Ref |
| --- | --- | --- | --- | --- | --- | --- |
| c.1360C>T | p.Leu454Phe | SCAR13 | severe to moderate intellectual disability; developmental delay; aggressive behaviour; disordered speech; cerebellar atrophy; gait and stance ataxia; seizure; nystagmus | Loss of function; missense mutation | N/A | [1] |
| c.2652_2654del and c.2660+2T>G | p.Asn885del + alternate splicing | SCAR13 | Mild to severe intellectual disability; global developmental delay; impaired or absent speech; progressive cerebellar atrophy; stance ataxia; gait ataxia; dysarthria; tremors; hyperreflexia; mild pyramidal signs | Loss of function; non-functional receptors lacking 7TM domain and/or abnormal or absent C-tail | N/A | [2] |
| c.785A>G | p.Tyr262Cys | SCA44 | No cognitive impairment; late onset cerebellar ataxia; spasticity | Gain of function; missense mutation | enhanced activity in luciferase reporter assay | [3] |
| c.2375A>G | p.Tyr792Cys | SCA44 | No cognitive impairment; late onset cerebellar ataxia; spasticity | Gain of function; missense mutation | enhanced activity in luciferase reporter assay | [3] |
| c.3165dup | p.Gly1056Argfs*49 | SCA44 | intellectual disability; cerebellar ataxia without cerebellar atrophy; normal brain imaging | Loss of function; modified and truncated C-tail | Loss of mGlu<sub>1</sub>- Homer clustering; reduced pERK1/2; reduced activity in luciferase reporter assay | [3] |

N/A; no functional experiments have been carried out for these variants
[1] Davarniya et al., 2015; [2] Guergueltcheva et al., 2012; [3] Watson et al., 2017

To assess allosteric modulator pharmacological profiles at SCA-associated mGlu_1_ mutants, two structurally and functionally distinct PAMs were chosen for loss-of-function mutants; Ro 67-4853 and CPPHA. Ro 67-4853 was one of the first described selective mGlu_1_ PAMs, whereas CPPHA is a dual mGlu_1_/mGlu_5_ PAM proposed to bind to secondary allosteric site ^28–32^. For gain-of-function mutants, JNJ16259685 and VU0469650 were chosen as structurally distinct, potent CNS penetrant NAMs ^33–35^. JNJ16259685 is efficacious in preclinical models of spinocerebellar ataxia without mGlu_1_ ^24^. Additionally, the FDA-approved anti-helminthic drug nitazoxanide was included as nitazoxanide was identified as a putative mGlu_1_ NAM from *in silico* drug screening, and later shown to reduce activity of both Y262C and Y792C mutations in reporter gene assays ^14,27^. However, initial screening in the current study revealed nitazoxanide displayed non-selective effects on cellular calcium signalling, inhibiting iCa^2+^ mobilisation induced by purinergic and muscarinic agonists in non-transfected HEK293A cells, as well as mGlu_1_ induced iCa^2+^ mobilisation (Figure S1). Given the non-selective effects of nitazoxanide, it was not carried forward into testing for modulation of mutant mGlu_1_ function.

### 3.1. Initial characterisation of tetracycline inducible mGlu_1_ cell lines

Initially, iCa^2+^ mobilisation assays were carried out for glutamate and quisqualate at WT and mutant mGlu_1_ receptors at different expression levels induced by increasing tetracycline concentrations (Figure S2). No orthosteric agonist-mediated iCa^2+^ mobilisation was evident under any tested tetracycline concentrations for G1056Rfs*49. With the exception of G1056Rfs*49, all variants displayed a biphasic receptor titration profile for both agonists, with an increasing E_max_ from 10–30ng/ml of tetracycline and a subsequent decrease in E_max_ at concentrations above 30ng/ml of tetracycline for both agonists. For the majority of constructs there was also considerable mGlu_1_ mediated iCa^2+^ mobilisation in the absence of tetracycline for both agonists, the exception being L454F. Given the maximal response to both agonists were observed after 30ng/ml tetracycline induction for WT and most mutants, these conditions were chosen for all subsequent functional and expression assays.

### 3.2. G1056Rfs*49 results in loss of mGlu_1_ expression

Parallel to functional assays, cell surface expression of mutant receptors was compared to WT mGlu_1_ using flow cytometry (Figure 1). A fluorescent anti-cMyc antibody was used to detect surface expression via binding to the N-terminal cMyc tag fused to mGlu_1_ constructs. No difference in expression was observed under baseline conditions (Table S2). Tetracycline (30ng/ml) resulted in increased receptor expression relative to non-transfected control cells (Figure 1; Table S2). Interestingly, tetracycline (1µg/ml) resulted in a further increase in receptor expression relative to 30ng/ml, indicating the loss of iCa^2+^ functional responses at higher tetracycline concentrations are not due to a reduction in expression. With the exception of G1056Rfs*49, no mutants significantly affected mGlu_1_ surface expression at 30ng/ml tetracycline. Expression of G1056Rfs*49 was significantly lower (6.9-fold) compared to WT under these conditions (Figure 1). FACS was carried also out on permeabilised G1056Rfs*49 cells to determine if loss of expression was cell surface specific, or if mGlu_1_ was being expressed intracellularly. Expression of G1056Rfs*49 was significantly different to WT under permeabilised conditions (Figure 1).

### 3.3. mGlu_1_ mutations differentially effect orthosteric agonist potency, affinity and efficacy

Potency and E_max_ estimates derived from iCa^2+^ mobilisation experiments for both agonists after induction with 30ng/ml tetracycline revealed quisqualate was significantly more potent than glutamate for all mGlu_1_ variants (Figure 2, Table S3). There were no differences in E_max_ estimates between agonists with the exception of N885del, where quisqualate E_max_ was reduced 2-fold relative to glutamate. Compared to WT, glutamate and quisqualate potencies at Y262C and Y792C were similar in iCa^2+^ mobilisation assays. However, glutamate potency was significantly reduced by 6.3-fold at L454F, with no significant change in quisqualate potency, compared to WT. The potency of both glutamate and quisqualate were significantly reduced for N885del, by 5.5-fold and 6.4-fold, respectively. Maximal responses were similar between WT, L454F, Y262C and Y792C for both agonists. However, quisqualate E_max_ estimates were significantly reduced by 1.5-fold for N885del compared to WT (Figure 2, Table S3).

**Figure 2.**
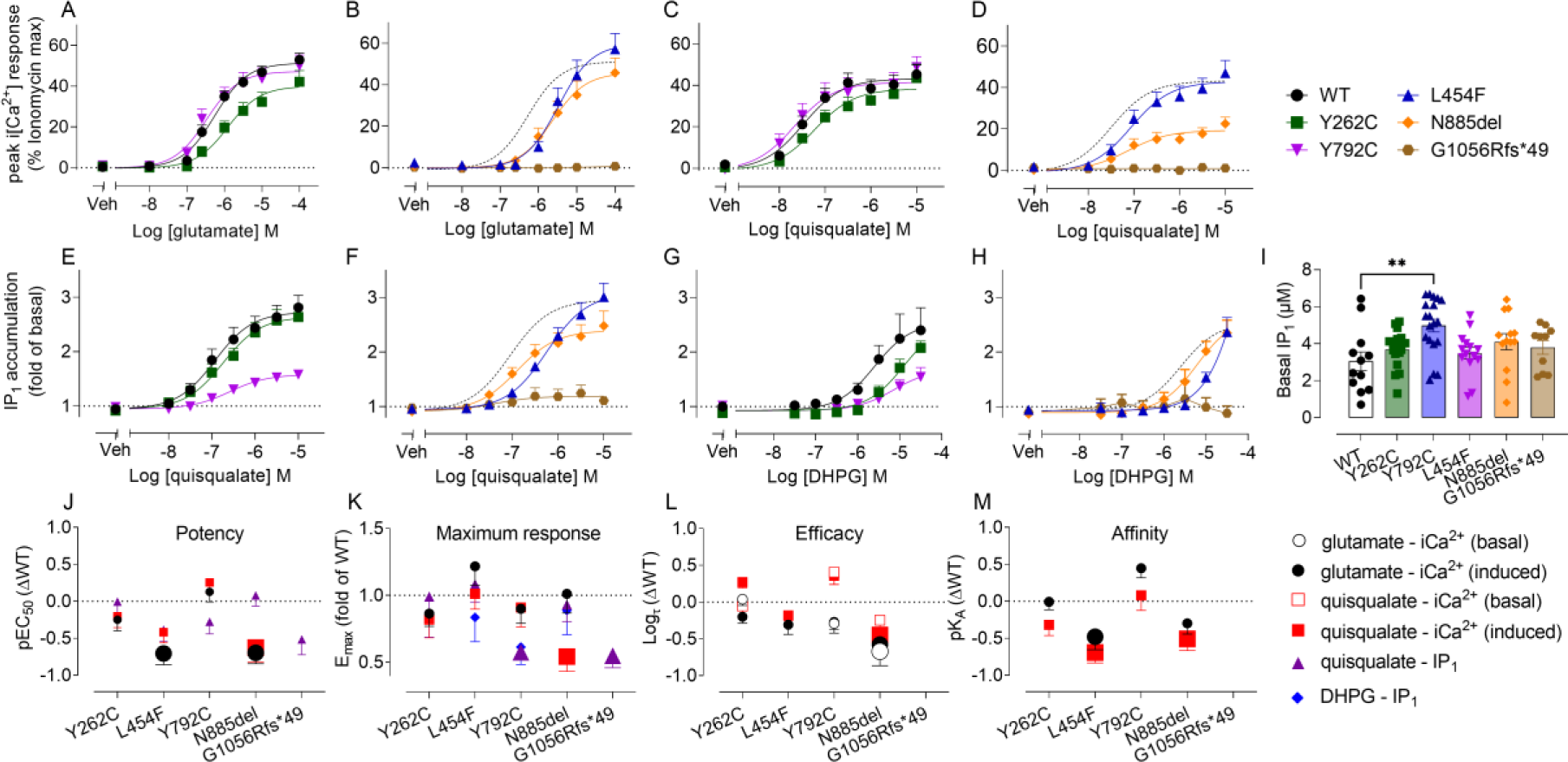
mGlu_1_ mutations differentially influence orthosteric ligand pharmacology in iCa^2+^ mobilisation and IP_1_ accumulation in Flp-In TRex HEK293 cells expressing mutant and wild type mGlu_1_ receptors. Concentration response curves for iCa^2+^ mobilisation assays to the orthosteric agonists glutamate (A-B) and quisqualate (C-D) and in IP_1_ accumulation for quisqualate (E-F) and DHPG (G-H) following overnight induction with 30ng/ml tetracycline. (I) Y792C significantly increases basal IP_1_ accumulation relative to WT. (J-M) Quantification of orthosteric agonist potency, maximum response, efficacy and affinity differences between WT and mutant mGlu_1._ Oversized symbols denote estimates are significantly different to WT. All IP_1_ accumulation assays were performed in the presence of 10U/mL glutamic-pyruvic transaminase (GPT) to minimise the influence of ambient glutamate. Data are expressed as mean + S.E.M of at least 5 experiments performed in duplicate (refer to Tables S3 & S4 for n numbers for each experiment). Error bars not shown lie within the dimensions of the symbols. Significance was determined using an ordinary one-way ANOVA test with Dunnett’s (for basal IP_1_) or Sidak’s (for pharmacological parameters) post-hoc tests.

Concentration-response experiments carried out at a single receptor expression level only provide potency estimates, which represent a mixture of ligand affinity and efficacy. The benefit of inducible cell lines is receptor expression can be titrated using different concentrations of tetracycline. By assaying agonist activity at different receptor levels and fitting a model of receptor depletion, global affinity (K_A_) estimates and efficacy (log_τ_) estimates under basal (no tetracycline) and induced (i.e. 30ng/ml tetracycline) conditions were derived for each ligand using global curve fitting across all tetracycline concentrations in iCa^2+^ mobilisation assays (Figure 2, Table S4). WT and all mutants displayed significantly higher affinity estimates for quisqualate relative to glutamate. Compared to WT, L454F had significantly lower affinity estimates for both glutamate (3-fold) and quisqualate (5-fold), with N885del also having a 3-fold lower affinity estimate for quisqualate. No other mutants significantly affected the affinity of either agonist. Receptor induction resulted in a significant increase in efficacy for both agonists for WT and all mutants relative to basal conditions (Table S4). Glutamate had significantly lower efficacy at N885del compared to WT under both basal (5-fold reduction) and induced (4-fold reduction) conditions (Figure 2). Similarly, quisqualate efficacy was significantly reduced by 2.8-fold for N885del under induced conditions.

### 3.4. mGlu_1_ mutations differentially effect distinct measures of canonical signalling

Transient (∼2 min) iCa^2+^ mobilisation assays were compared to another measure of canonical mGlu_1_-G_q_ signalling; extended (∼1 hr) IP_1_ accumulation assays (Figure 2). Given the extended nature of IP_1_ accumulation assays and the potential confound of glutamate being produced and released by HEK293 cells over time, GPT was added to IP_1_ accumulation assay buffer to eliminate ambient glutamate. This precluded the use of glutamate as an orthosteric ligand in these assays. Instead, the synthetic group I mGlu selective surrogate agonist DHPG was added as a second agonist in IP_1_ accumulation assays. Potency (pEC_50_) and E_max_ values were derived to determine differences between mutants and between functional signalling outcomes (Table S3).

In IP_1_ accumulation assays, Y792C displayed an increase in basal IP_1_ levels relative to WT, indicative of enhanced constitutive activity (Figure 2). In contrast to iCa^2+^ mobilisation, G1056Rfs*49 had a small but detectable response to quisqualate in IP_1_ accumulation assays. No mutation significantly affected quisqualate potency compared to WT (Table S3). Only Y792C and G1056Rfs*49 significantly affected quisqualate E_max_ estimates, with a 1.7-fold and 1.8-fold reduction compared to WT, respectively (Figure 2, Table S3). When comparing quisqualate between iCa^2+^ mobilisation and IP_1_ accumulation assays, potency estimates were similar for all mutants with the exception of Y792C, which had 14.4-fold decreased potency for IP_1_ accumulation relative to iCa^2+^ mobilisation. In contrast to quisqualate, G1056Rfs*49 had no response to DHPG in IP_1_ accumulation assays (Figure 2). DHPG curves were right shifted relative to WT for all other mutations, prohibiting derivation of potency estimates due to solubility limits for concentrations above 30 µM (Figure 2). However, a comparison of maximum response values at 30 µM revealed no significant differences between WT and any mutant (Figure 2, Table S3).

### 3.5. PAMs display altered pharmacology at mGlu_1_ loss-of-function mutations

To characterise the impact of PAMs on wild type and loss-of-function mutant mGlu_1_, a titration paradigm was utilised, in which different concentrations of PAMs were applied in the absence and presence of EC_20_ of orthosteric agonists to investigate both intrinsic agonism and modulatory effects. Potency and maximal effect parameters for PAMs were derived in the absence (pEC_50_/E_max_) and presence (pMod_50_/Mod_max_) of agonist. No response was observed to either PAM when applied alone in iCa^2+^ mobilisation assays for WT or mutant receptors (Figure 3), indicating Ro 67-4853 and CPPHA lack intrinsic agonist efficacy at mGlu_1_. Both Ro 67-4853 and CPPHA enhanced iCa^2+^ mobilisation induced by EC_20_ glutamate or quisqualate in a concentration dependent manner at WT mGlu_1_ (Figure 3). However, CPPHA concentration response curves did not reach a plateau, prohibiting derivation of pEC_50_ and pMod_50_ values. As such, Mod_max_ was defined as the span between the baseline EC_20_ response and the response at the highest concentration of PAM for CPPHA (Table S5).

**Figure 3.**
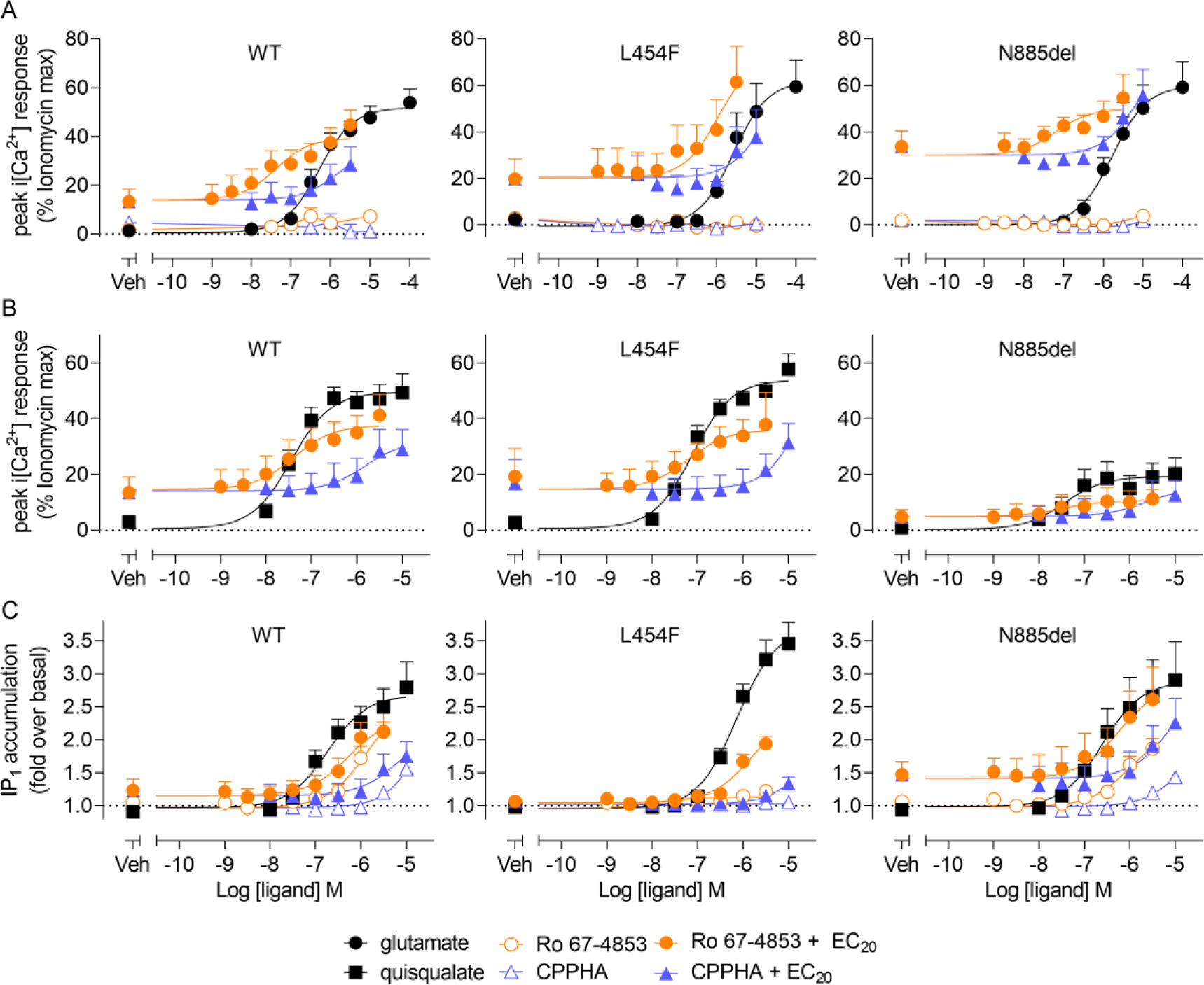
Loss-of function mGlu_1_ mutations differentially affect PAM intrinsic agonism and modulation of glutamate and quisqualate induced iCa^2+^ mobilisation and IP_1_ accumulation in Flp-In TRex HEK293 cells. (A) Concentration response curves for PAM intrinsic agonism (open symbols) and modulation of an EC_20_ concentration of glutamate (closed symbols) in iCa^2+^ mobilisation. (B) Concentration response curves for PAM modulation of an EC_20_ concentration of quisqualate in iCa^2+^ mobilisation. (C) Concentration response curves for PAM intrinsic agonism (open symbols) and modulation of an EC_20_ concentration of quisqualate (closed symbols) in IP_1_ accumulation. IP_1_ accumulation assays were performed in the presence of 10U/mL GPT to minimise the contribution of ambient glutamate. Data represent mean + SEM of at least 3 experiments performed in duplicate (refer to Table S6 for exact n numbers for each dataset). Error bars not shown lie within the dimensions of the symbol

Potency and maximum effect estimates for Ro 67-4853 and CPPHA were not different between modulation of glutamate and quisqualate for WT mGlu_1_ (Figure 3, Figure 6). Ro 67-4853 curves could not be fitted for modulation of glutamate at L454F or for modulation of quisqualate at N885del mutants, indicating a loss of Ro 67-4853 potentiator potency relative to WT (Figure 3). For Ro 67-4853 modulation of glutamate at N885del and quisqualate at L454F, there were no differences in pMod_50_ or Mod_max_ estimates, relative to WT (Figure 6, Table S5). Similar to WT, CPPHA modulation curves did not reach a plateau for either agonist at L454F or N885del. However, neither mutant significantly affected Mod_max_ estimates for CPPHA for modulation of either glutamate or quisqualate induced iCa^2+^ mobilisation (Figure 6, Table S5).

Unlike iCa^2+^ mobilisation, PAMs exhibited intrinsic agonism in IP_1_ accumulation assays for WT and both loss-of-function mutant variants (Figure 3C). A lack of plateau again precluded derivation of potency estimates, but comparison of maximum response at the highest concentration tested revealed no significant differences in E_max_ for either PAM alone when comparing N885del and WT (Figure 6, Table S5). L454F significantly reduced E_max_ of Ro 67-4853 compared to WT, with no detectable CPPHA agonism. Despite reduced agonist activity, Ro 67-4853 modulated quisqualate induced IP_1_ accumulation at L454F, with similar pMod_50_ and Mod_max_ values compared to WT (Figure 6, Table S5). Interestingly, pMod_50_ values for Ro67-4853 modulation of quisqualate were significantly lower in IP_1_ accumulation compared to iCa^2+^ mobilisation for both WT (16-fold) and L454F (14.8-fold) (Figure 6, Table S5). Ro 67-4853 and CPPHA modulation curves failed to reach a plateau for modulation of quisqualate for N885del, but Mod_max_ estimates were comparable to WT (Table S5).

Similar to orthosteric agonist potency estimates, potency of allosteric modulators represents a composite of pharmacological properties. Allosteric ligand pharmacology is primarily defined by two main properties: affinity and cooperativity. We used an operational model of allosterism to derive affinity (pK_B_) and cooperativity (logβ) estimates for PAM modulation of orthosteric agonist responses in both iCa^2+^ mobilisation and IP_1_ accumulation assays (Table S5). Concentration response curves that did not reach plateau could not be fit, and as such only Ro 67-4853 estimates were derived for certain agonist/mutant combinations. pK_B_ and logβ estimates for Ro 67-4853 were similar between modulation of glutamate and quisqualate for WT mGlu_1_ in iCa^2+^ mobilisation. Similarly, Ro 67-4853 pK_B_ and logβ estimates were not significantly different between WT and L454F for modulation of quisqualate, and between WT and N885del for modulation of glutamate in iCa^2+^ mobilisation (Figure 6, Table S5). However, when comparing between assays, Ro 67-4853 apparent affinity at WT and L454F was significantly lower (∼20-25 fold) in IP_1_ accumulation compared to iCa^2+^ mobilisation (Figure 6, Table S5). Ro 67-4853 cooperativity was best fit as neutral at WT (as determined by an extra sum-of-squares *F*-test), suggesting that a majority of PAM activity could be attributed to intrinsic agonism. Cooperativity estimates for Ro 67-4853 with quisqualate were not significantly different between IP_1_ accumulation and iCa^2+^ mobilisation for L454F (Table S5).

### 3.6. Ro 67-4853 can rescue glutamate-mediated iCa^2+^ mobilisation at both L454F and N885del

Despite a loss of potency for Ro 67-4853 modulation of glutamate induced iCa^2+^ mobilisation, both PAMs modulated EC_20_ glutamate at high concentrations at both L454F and N885del. Given that glutamate displayed a significant loss of potency at both L454F and N88del in iCa^2+^ mobilisation assays relative to WT, we next tested whether a single high concentration of either PAM could restore glutamate potency to WT levels (Figure 4). Ro 67-4853 and CPPHA (10µM) left-shifted L454F glutamate concentration response curves by 2-3-fold compared to vehicle. The resulting glutamate potency estimate in the presence of Ro 67-4853 was no longer significantly different from potency estimates at WT (Table 2). At N885del, glutamate potency was significantly shifted by both Ro 67-4853 (5-fold) and CPPHA (3.2-fold) and were no longer significantly different to WT potency estimates (Figure 4, Table 2). A slight increase in glutamate E_max_ also evident in the presence of both PAMs at L454F, but this was not significantly different to vehicle E_max_, or WT estimates (Figure 4).

**Figure 4.**
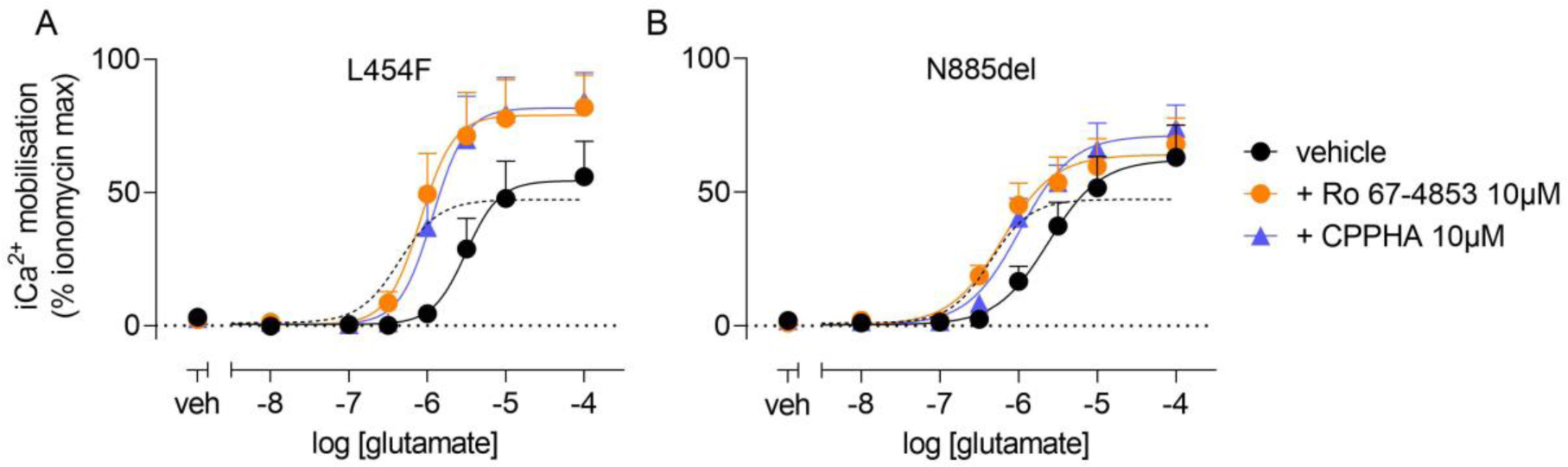
Ro 67-4853 and CPPHA can rescue glutamate responses at loss-of function mGlu_1_ mutations in Flp-In TRex HEK293 cells. Concentration response curves for glutamate in the absence or presence of 10µM Ro 67-4853 or CPPHA at L454F (A) and N885del (B). Both PAMs increased glutamate potency and E_max_ relative to vehicle treatment at L454F, and increased potency at N885del. WT glutamate responses are shown in dashed lines for reference. Data represent mean + SEM of 5 experiments performed in duplicate.

**Table 2.**
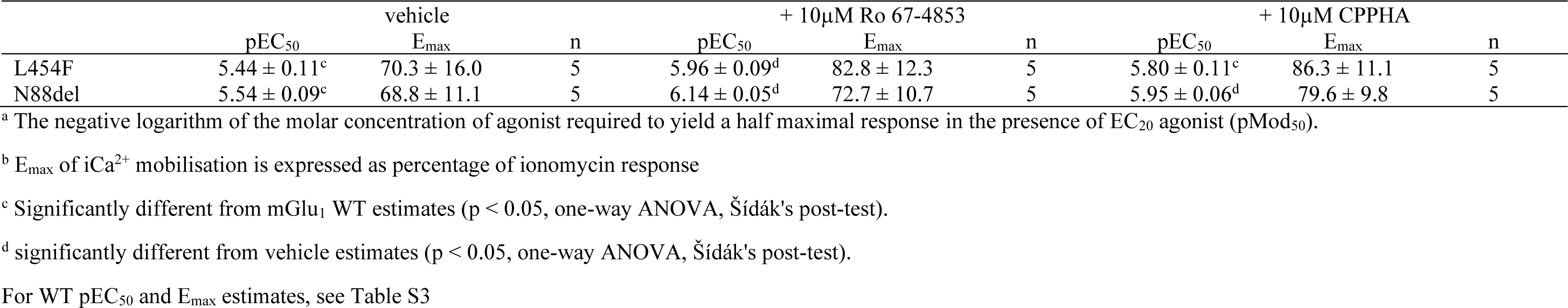
Potency and maximum response estimates for modulation of glutamate induced iCa^2+^ mobilisation by a single concentration of PAMs in Flp-In TRex HEK293 sexpressing loss-of-function mutations. Data are mean ± SEM of indicated number (n) of independent experiments performed in duplicate.

| | vehicle | | | + 10μM Ro 67-4853 | | | + 10μM CPPHA | | |
| --- | --- | --- | --- | --- | --- | --- | --- | --- | --- |
|  | pEC<sub>50</sub> | E<sub>max</sub> | n | pEC<sub>50</sub> | E<sub>max</sub> | n | pEC<sub>50</sub> | E<sub>max</sub> | n |
| L454F | 5.44 ± 0.11<sup>c</sup> | 70.3 ± 16.0 | 5 | 5.96 ± 0.09<sup>d</sup> | 82.8 ± 12.3 | 5 | 5.80 ± 0.11<sup>c</sup> | 86.3 ± 11.1 | 5 |
| N88del | 5.54 ± 0.09<sup>c</sup> | 68.8 ± 11.1 | 5 | 6.14 ± 0.05<sup>d</sup> | 72.7 ± 10.7 | 5 | 5.95 ± 0.06<sup>d</sup> | 79.6 ± 9.8 | 5 |

<sup>a</sup> The negative logarithm of the molar concentration of agonist required to yield a half maximal response in the presence of EC<sub>20</sub> agonist (pMod<sub>50</sub>).
<sup>b</sup> E<sub>max</sub> of $iCa^{2+}$ mobilisation is expressed as percentage of ionomycin response
<sup>c</sup> Significantly different from mGlu<sub>1</sub> WT estimates (p < 0.05, one-way ANOVA, Šidák's post-test).
<sup>d</sup> significantly different from vehicle estimates (p < 0.05, one-way ANOVA, Šidák's post-test).
For WT pEC<sub>50</sub> and E<sub>max</sub> estimates, see Table S3

### 3.7. Ro 67-4853 has enhanced agonist activity in IP_1_ accumulation at gain-of-function mGlu_1_ mutations

PAM agonism was initially tested at both Y262C and Y792C, to determine if putative gain-of-function manifested as enhanced PAM activity. Neither PAM had agonist activity at either mutant in iCa^2+^ mobilisation assays (Figure 5). However, in IP_1_ accumulation assays, Ro 67-4853 displayed enhanced intrinsic agonism relative to WT (Figure 5C). Both Y262C and Y792C increased Ro 67-4853 potency as, at WT, Ro 67-4853 pEC_50_ values could not be derived, but both mutants had full concentration response curves and resulting potency estimates were similar between mutants (Figure 5C, Figure 6, Table S6). Ro 67-4853 E_max_ estimates for both mutants were similar to WT (Figure 6, Table S6). Interestingly, CPPHA lost agonist activity at Y262C, but retained function similar to WT at Y792C (Figure 5C).

**Figure 5.**
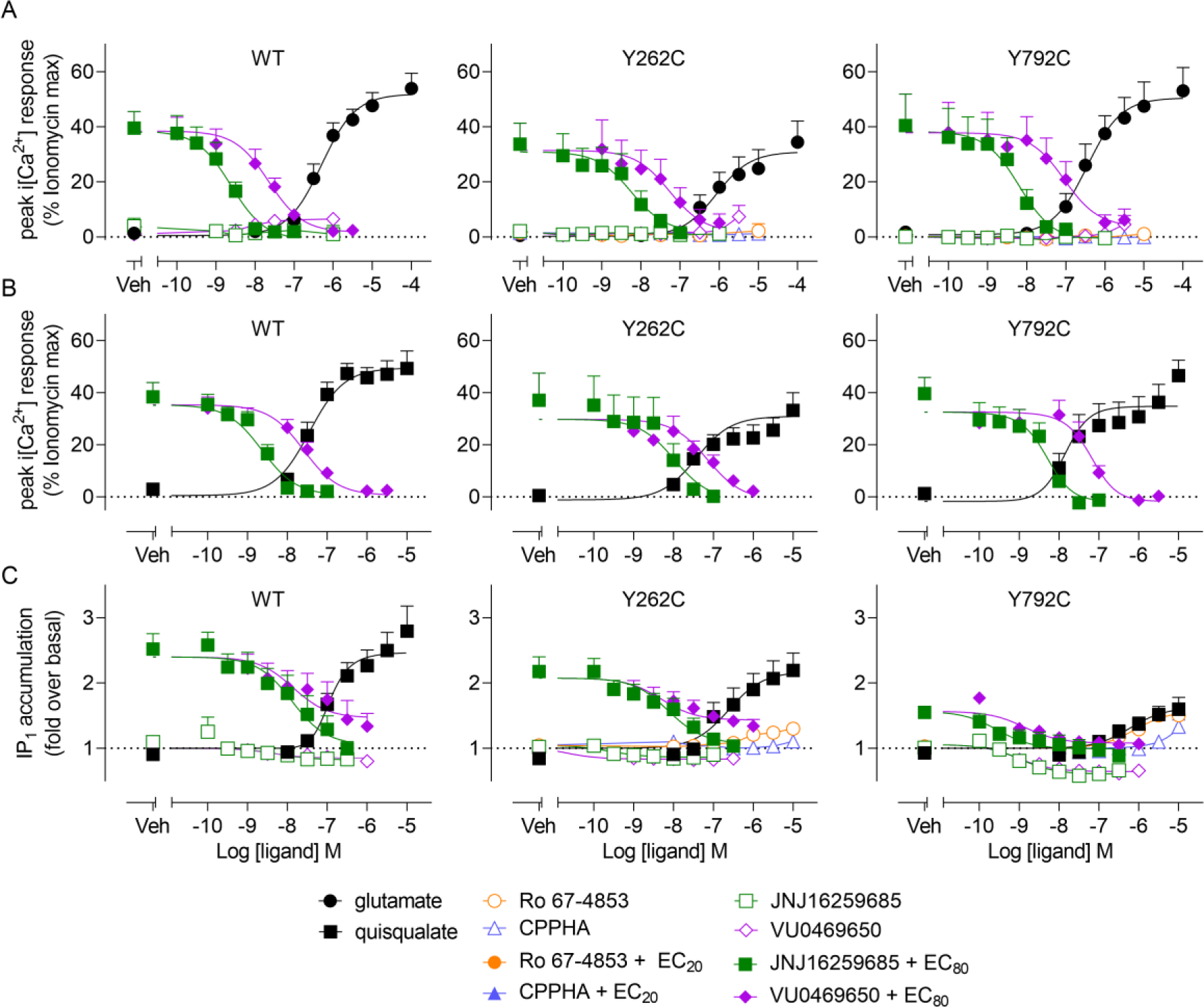
Gain-of function mGlu_1_ mutations differentially affect PAM intrinsic agonism, NAM inverse agonism and modulation of glutamate and quisqualate induced iCa^2+^ mobilisation and IP_1_ accumulation in Flp-In TRex HEK293 cells. (A) Concentration response curves for PAM and NAM agonism/inverse agonism (open symbols) and modulation of an EC_80_ concentration of glutamate (closed symbols) in iCa^2+^ mobilisation. (B) Concentration response curves for NAM modulation of an EC_80_ concentration of quisqualate in iCa^2+^ mobilisation. (C) Concentration response curves for PAM and NAM agonism/inverse agonism (open symbols) and NAM modulation of an EC_80_ concentration of quisqualate (closed symbols) in IP_1_ accumulation. IP_1_ accumulation assays were performed in the presence of 10U/mL GPT to minimise the contribution of ambient glutamate. Data represent mean + SEM of at least 3 experiments performed in duplicate (refer to Tables S3 & S5 for n numbers for each experiment). Error bars not shown lie within the dimensions of the symbol.

### 3.8. mGlu_1_ NAMs reduce constitutive activity at Y792C in IP_1_ accumulation

To characterise the inhibitory effects of NAMs on mutant and WT mGlu_1_, VU0469650 and JNJ16259685 were tested in the absence and presence of EC_80_ orthosteric agonist. In iCa^2+^ mobilisation assays, neither NAM had inverse agonist activity when applied alone, but both JNJ16259685 and VU0469650 fully inhibited glutamate and quisqualate EC_80_ responses at WT, L454F and N885del (Figure 5A & B). Comparison of pIC_50_ and inhibitory I_max_ values revealed no significant differences for modulation of either agonist by JNJ16259685 and VU0469650 at either mutant compared to WT, nor any differences between the two agonists within each mutant (Figure 6, Table S6).

**Figure 6.**
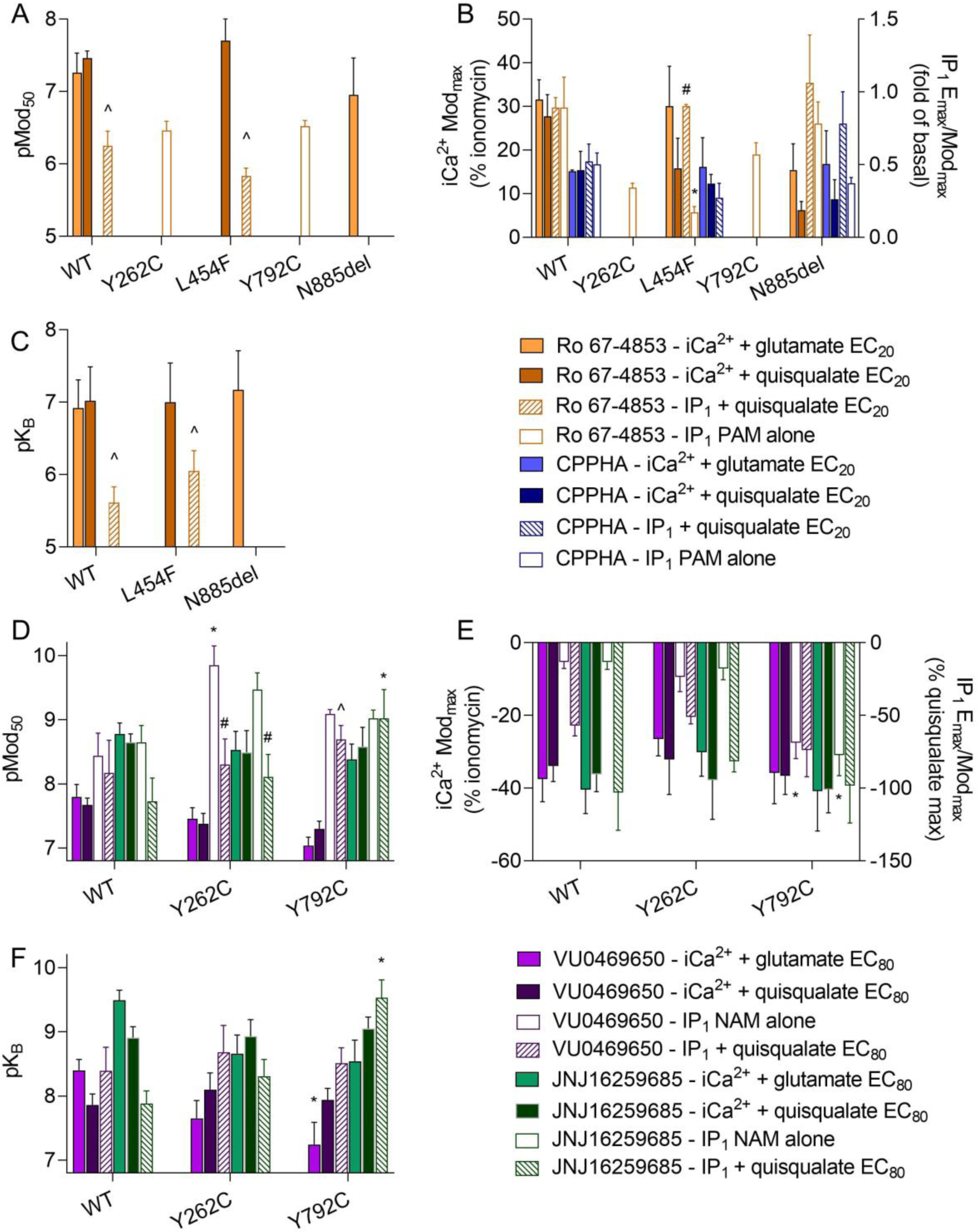
Quantification of potency, affinity and maximum response parameters for PAM and NAM agonism and modulation glutamate and quisqualate induced iCa^2+^ mobilisation and IP_1_ accumulation in Flp-In TRex HEK293 cells expressing WT and mutant mGlu_1_. pEC_50_/pMod_50_ and E_max_/Mod_max_ are derived from fitting concentration response curves for allosteric modulators in the absence or presence of EC_20_ (PAMs) or EC_80_ (NAMs) of glutamate or quisqualate to a variable four-parameter logistic equation. pK_B_ is derived from fitting the curves described above to an operational model of allosterism. Mod_max_ of iCa^2+^ mobilisation is the span between EC_20_/EC_80_ baseline and the top or bottom plateau of PAM or NAM curves, expressed as percentage of ionomycin response. E_max_/Mod_max_ of IP_1_ accumulation for PAMs is the span between baseline (E_max_) or baseline in the presence of EC_20_ (Mod_max_) and responses at highest concentration of PAM and are expressed as fold of basal. Due to difference in signal window for Y792C compared to WT and Y262C, Mod_max_ of IP_1_ accumulation for NAMs is presented the span between EC_80_ baseline and the bottom plateau of NAM curves, expressed as percentage of quisqualate response. * denotes significantly different from mGlu1 WT response. ^ denotes significantly different from modulation of quisqualate mediated iCa mobilisation. # denotes significantly different from PAM alone in IP1 modulation. Data are expressed as mean + S.E.M of at least 5 experiments performed in duplicate. Error bars not shown lie within the dimensions of the symbols. Significance was determined using an ordinary one-way ANOVA test with Sidak’s post-hoc test.

In IP_1_ accumulation assays, both NAMs displayed inverse agonism at WT and both mutants (Figure 5C). Both JNJ16259685 and VU0469650 had enhanced inverse agonist activity at Y792C, with significantly lower I_max_ estimates compared to WT (Figure 6, Table S6). JNJ16259685 fully inhibited the IP_1_ accumulation induced by EC_80_ quisqualate for WT and both mutants. JNJ16259685 was significantly more potent (∼19 fold) for modulation of quisqualate in IP_1_ accumulation at Y792C compared to WT (Figure 6, Table S6). VU0469650 fully inhibited quisqualate responses at Y792C, but only partially inhibited quisqualate responses at WT and Y262C (Figure 5C). Additionally, the pMod_50_ estimate for VU0469650 modulation of quisqualate in IP_1_ assays was significantly higher (∼8 fold) than Ca^2+^ mobilisation assays for Y792C (Figure 6, Table S6). At Y262C, intrinsic inverse agonist potency was significantly higher (10-35-fold) compared to modulatory potency for both NAMs (Figure 6, Table S6).

As for PAMs at loss-of-function mutants, we next used an operational model of allosterism to derive affinity and cooperativity estimates for NAM modulation of orthosteric agonist responses (Figure 6, Table S6). As JNJ16259685 fully inhibited orthosteric agonists in both iCa^2+^ mobilisation and IP_1_ accumulation assays, and VU0469650 acted as full NAM in iCa^2+^ mobilisation assays, cooperativity estimates could only be derived for VU0469650 in IP_1_ accumulation. VU0469650 affinity for modulation of glutamate in iCa^2+^ mobilisation was significantly reduced by 14-fold at Y792C compared to WT (Figure 6, Table S6). Conversely, JNJ16259685 affinity for modulation of quisqualate in IP_1_ accumulation was significantly increased by 54-fold, compared to WT (Figure 6, Table S6). Comparison of cooperativity estimates for VU0469650 in IP_1_ accumulation between WT and Y262C revealed no significant differences.

## 4. Discussion

Rare genetic forms of SCA are associated with mGlu_1_ mutations in preliminary clinical reports (Cabet et al. 2019; Davarniya et al., 2015; Guergueltcheva et al., 2012; Ngo et al., 2020; Protasova et al., 2023; Rasool et al., 2021; Watson et al., 2017; Yousaf et al., 2022). Although the clinical phenotypes and location of mutations within mGlu_1_ have been identified, how these mutations result in abnormal mGlu_1_ function, and how this then contributes to SCA progression, is unknown. Here, we aimed to characterise wild type mGlu_1_ function and compare to mGlu_1_ harbouring disease-associated mutations using expression analysis, iCa^2+^ mobilisation and IP_1_ accumulation assays. Characterisation included both orthosteric agonists and allosteric ligands to determine the feasibility of restoring aberrant function using small molecules. Expression analysis revealed G1056Rfs*49 leads to an almost complete loss of mGlu_1_ expression. L454F and N885del mutations caused a loss of function through changes in orthosteric agonist affinity and efficacy. The gain-of-function mutation Y792C enhanced constitutive activity which resulted in reduced orthosteric agonist responses. Loss-of-function mutations also affected PAM pharmacology, whereas NAMs mostly retained activity at gain-of-function mutations. Finally, a putative mGlu_1_ NAM, nitazoxanide, was revealed to exert non-selective effects on cellular iCa^2+^ signalling. Overall, this study provides critical insights into the mechanisms by which naturally occurring mutations can impact of mGlu_1_ function and closes the gap between cellular outcomes and the pathology of hereditary ataxias.

In the current study, the G1056Rfs*49 mutation resulted in a significant loss in expression of mGlu_1_ relative to wild type. The C-terminal domain that is lost upon the frame-shift contains binding sites for multiple protein partners, including cytoskeletal and scaffolding proteins like β-tubulin, tamalin and Homer1/2b (see Figure 1) ^36–41^. Homer binds to proline-rich sections in group I mGlu C-termini and is critical for localising mGlu_1_ to discrete sites at postsynaptic excitatory synapses and linking mGlu_1_ and mGlu_5_ to iCa^2+^ mobilisation via direct interaction with IP_3_ receptors ^36,40,41^. Indeed, the G1056Rfs*49 mutation results in a complete ablation of clustering with Homer 2b in recombinant cells (Watson et al., 2017). The lack of cell surface expression and subsequent dramatic reduction in functional responses in the current study likely results from this poor Homer clustering, as both Homer 1 and 2b are endogenously expressed in HEK293A cells (data available from https://www.proteinatlas.org/ENSG00000103942-HOMER2 and https://www.proteinatlas.org/ENSG00000152413-HOMER1)^42^. However, a small functional response was observed in IP_1_ accumulation assays with quisqualate, but not DHPG. Both quisqualate and DHPG are membrane impermeable agonists, but quisqualate can be actively transported across the membrane, suggesting the small functional responses evident in IP_1_ accumulation assays results from activation of intracellular receptors ^43^. Activation of a small intracellular population of receptors is further supported by evidence that non-clustered G1056Rfs*49 receptors had small but detectable activity in both ERK1/2 and a reporter gene assays, both of which were significantly reduced compared to WT^14^. The mechanism by which G1056Rfs*49 results in loss of mGlu_1_ function suggests modulation by mGlu_1_-targeting pharmacological agents would not be viable.

L454F and N885del have been suggested as loss-of-function mutations without supporting pharmacological evidence^12,13^. In the current study, receptor titration analysis revealed L454F primarily reduced orthosteric ligand affinity, whereas N885del affected glutamate efficacy and both quisqualate affinity and efficacy, demonstrating multiple mechanisms through which loss-of-function can occur. These different mechanisms are perhaps not surprising when the locations of the mutated residues are considered (Figure 1). L454 is located in the extracellular ligand binding domain, where a bulky phenylalanine would be predicted to interfere with pocket formation and subsequent ligand binding^13^. Conversely, N885 is located in the C-terminal tail and may be involved in interactions between mGlu_1_ and intracellular effectors and scaffolding proteins. While N885del co-occurs in patients with an intron 8 splicing mutation, and the resulting alternate splicing and truncated/altered C-terminus likely dominates subsequent effects on mGlu_1_ function, it is interesting a single amino acid deletion also has functional consequences ^12^. The proximal C-terminal region containing N885 is critical for interactions between mGlu_1_ and other receptors (NMDA and adenosine receptors), and ERK1/2 also binds between K841 and N885 to regulate mGlu_1_ expression and canonical signalling ^12,44–46^ Importantly, N885del did not affect IP_1_ signalling by either orthosteric agonists or PAMs, indicating a calcium-specific affect of this mutation. Indeed, N885 is also predicted to be in a putative calmodulin binding domain, which may account for selective affects on Ca^2+^ signalling^47^. Taken together these data reiterate the need for extensive functional characterisation of receptor function across multiple assays, using pertinent pharmacological parameters, to fully appreciate the complexity of the effects of naturally mutations on mGlu_1_ signalling.

Neither putative gain-of-function mutation affected orthosteric agonist affinity or efficacy, indicating more subtle effects on mGlu_1_ function. Y792C increased constitutive mGlu_1,_ activity which was detectable in IP_1_ accumulation but not iCa^2+^ mobilisation. Increased constitutive activity likely leads to run-down of canonical signalling pathways upon extended receptor activation, manifesting as reduced agonist response in IP_1_ accumulation assays. Such run-down could occur through depletion of signalling molecules, as second messengers have been suggested as limiting factors in receptor over-expression systems^48^. Alternatively, increased basal activity may result in higher levels of receptor desensitisation, as group I receptors undergo rapid desensitisation upon activation ^49–53^. Y262C had a different profile to Y792C, with no changes in canonical receptor signalling, despite displaying similarly enhanced activity to Y792C in reporter gene assays in initial reports ^14^. Such assays required much longer incubation times than either of the functional endpoints measured here, indicating functional effects of Y262C may rely on extended agonist exposure. Indeed, the lack of GPT in buffers in prior studies suggest ambient glutamate was likely chronically activating mGlu_1_, as levels of glutamate produced by cells can reach micromolar concentrations under non-GPT conditions, and incubation with an orthosteric agonist reduced Y262C activity to WT levels ^14,54^. Regardless of the mechanism of gain-of-function, enhanced mGlu_1_ signalling in patients harbouring either mutation would likely be toxic to cerebellar Purkinje neurons and result in progressive cerebellar atrophy. Calcium-mediated excitotoxic cascades are a feature of multiple SCA animal models and can be directly linked to dysregulated mGlu_1_ signalling in many cases, positioning mGlu_1_ as a target across SCA subtypes ^3,4,55,56^.

Both Y262 and Y792 are located within receptor domains critical for the large-scale conformational changes associated with mGlu_1_ activation (Figure 1). Y262 is located within the large mGlu_1_ extracellular ligand binding VFD. Molecular dynamic simulations have revealed Y262 contributes to interactions between the ligand binding LB2 domains of each protomer that facilitate mGlu_1_ dimerisation and stabilise an active state ^57^. Y792 is located in TM6 within the 7 transmembrane region, which is critical for forming a signalling-competent dimer conformation upon receptor activation ^58,59^. Comparisons of inactive and active mGlu_1_ structures reveal both Y262 and Y792C display significant movements upon receptor activation and dimer compaction, bringing them to close proximity with the opposing protomer (see Figure 1)^59^. Mutation to a cysteine likely stabilises a conformation of the VFD or TM domains that increases activity, by either changing interactions within each protomer or between protomers within the dimer. Y262C potentially reduces the energy required for mGlu_1_ activation, as enhanced interaction between the LB2 domains would mimic changes seen upon glutamate binding ^57^. Alternatively, Y792C would stabilise a constitutively active TM6 conformation, potentially through formation of a disulphide bond between the two protomers. Indeed, the closely related mGlu_5_ is constitutively active when an I791C mutation is introduced to TM6 to form a disulphide bond between protomers ^58^. Y792 on each protomer are oriented in the right direction to form a disulphide bond, and the distance between the two residues in the active mGlu_1_ and mGlu_5_ structures are compatible with disulphide formation^58,59^. Similar to the current study, constitutive mGlu_5_ activity could be inhibited with a NAM^58^, highlighting the potential utility of allosteric modulation as an approach to restore receptor function in gain-of-function mutations.

Allosteric modulation of mGlu_1_ has proven effective in animal models of polyglutamine SCAs, in which the NAM JNJ16259685 and the PAM Ro 0711401 reverse motor deficits in models of moderate and severe SCA1, respectively ^21,24^. Despite increases/decreases in mGlu_1_ expression in these models, animals still express functionally intact mGlu_1_, which are amenable to modulation; this may change upon the introduction of mutations. The only evidence for allosteric modulation of mutant mGlu_1_ function comes from the effects of the putative mGlu_1_ NAM nitazoxanide, which reduced signalling at gain-of-function mutants ^14,27^. Here we show nitazoxanide has non-selective effects on Ca^2+^ signalling in recombinant cells, likely linked to its identified effects on depletion of intracellular Ca^2+^ stores ^60^. However, validated selective mGlu_1_ NAMs inhibited mutant receptor function in the current study. VU0469650 fully inhibited iCa^2+^ signalling, with lower cooperativity for inhibition of IP_1_ accumulation, similar to recent reports at WT mGlu_1_ ^61^. VU0469650 and JNJ16259685 were more potent as inverse agonists than NAMs at Y262C, suggesting there is a concentration window in which NAMs could reduce baseline activity without compromising physiological glutamate signalling. For loss-of-function mutants Ro 67-4853, but not CPPHA, rescued glutamate signalling in iCa^2+^ mobilisation at both L454F and N885del. Ro 67-4853 and CPPHA have differential modulatory effects on glutamate; Ro 67-4853 modulates both glutamate affinity and efficacy, whereas CPPHA only modulates glutamate efficacy in iCa^2+^ mobilisation ^61^. This would explain the inability of CPPHA to fully rescue glutamate function at the affinity driven L454F loss-of-function mutant, whereas N885del responded to both PAMs due to their effects on glutamate efficacy. Ro 67-4853 has the added advantage of lacking intrinsic agonism at L454F. “Pure PAMs” lacking intrinsic agonism are advantageous as intrinsic glutamatergic tone can be modulated without direct receptor activation ^26^. However, mGlu_1_ signalling is highly pleiotropic, with activation of other pathways such as β-arrestin, ERK1/2 and cAMP in both recombinant and native systems ^29,30,62–64^. It remains to be seen if this lack of Ro 67-4853 agonism is apparent across all signalling pathways, or indeed if any of the PAMs or NAMs tested here have differential pharmacology across other disease relevant signalling pathways

Overall, characterisation of the functional effects of SCA-associated mutations in human mGlu_1_ has highlighted the diversity of effects such mutations can have on receptor function. For the first time, we investigated the effects of both loss- and gain-of-function SCA-associated mGlu_1_ mutations on the potency, efficacy and affinity of orthosteric ligands across multiple measures of disease-relevant canonical signalling. Additionally, the potential utility of using both positive and negative allosteric modulators to restore aberrant mGlu_1_ function was explored. Loss-of-function mutations affected mGlu_1_ function through multiple mechanisms, affecting mGlu_1_ cell surface expression and both orthosteric ligand efficacy and/or affinity in a mutant specific manner. The putative gain-of-function mutant Y262C exhibited similar functional profiles to WT mGlu_1_ at both iCa^2+^ mobilisation and IP_1_ accumulation assays and requires further investigation across other pathways activated by mGlu_1_ or in different kinetic contexts. Conversely, the gain-of-function Y792C mutation enhanced mGlu_1_ constitutive activity, elevating baseline IP_1_ levels and reducing agonist responses in IP_1_ assays. PAM and NAM activity was mostly conserved across mutants relative to WT. However, a loss of intrinsic PAM agonism was evident in IP_1_ accumulation assays for L454F. Encouragingly, the PAM Ro 67-4853 could rescue glutamate signalling in iCa^2+^ mobilisation at the loss-of-function L454F and N885del mutations, restoring glutamate potency to WT levels. Additionally, two structurally divergent NAMs had inhibitory effects on enhanced constitutive activity at Y792C. Taken together, these data indicate loss- and gain-of-function SCA-associated mGlu_1_ mutations that affect constitutive activity and agonist affinity and/or efficacy may be amenable to pharmacological modulation, and mGlu_1_ allosteric modulators represent a viable starting point for potential therapeutics for these rare SCA subtypes.

## Abbreviations

CNS: central nervous system
CPPHA: *N*-[4-chloro-2-[(1,3-dihydro-1,3-dioxo-2H-isoindol-2-yl)methyl]phenyl]-2-hydroxybenzamide
DHPG: (S)-3,5-dihydroxyphenylglycine
DMEM: Dulbecco’s modified Eagle’s medium
EC_20_: 20% effective concentration
EC_50_: 50% effective concentration
EC_80_: 80% effective concentration
ERK1/2: extracellular signal-regulated protein kinase 1/2
FBS: fetal bovine serum
GPCR: G protein-coupled receptor
HBSS: Hank’s Balanced Salt Solution
HEK293A: human embryonic kidney 293
iCa^2+^: intracellular calcium
IP_1_: inositol 1-phosphate
JNJ16259685: (3,4-dihydro-2H-pyrano[2,3-b]quinolin-7-yl)-(cis-4-methoxycyclohexyl)-methanone
mGlu: metabotropic glutamate receptor
NAM: negative allosteric modulator
PAM: positive allosteric modulator
Ro 67-4853: (9H-xanthen-9-ylcarbonyl)-carbamic acid butyl ester
VU0469650: 3-[(3R)-3-methyl-4-(tricyclo[3.3.1.13,7] dec-1-ylcarbonyl)-1-piperazinyl]-2-pyridinecarbonitrile

## Acknowledgements

This work was supported by a National Ataxia Foundation Early Career Investigator Award to SDH. Research on metabotropic glutamate receptor allosteric modulators within the Endocrine and Neuropharmacology lab was supported by an Australian Research Council Future Fellowship (FT170100392) and a National Health and Medical Research Council Australia Ideas grant (2002947) awarded to KJG.

## Authorship Contributions

Participated in research design: Wang, Hellyer, Gregory; conducted experiments: Wang, Muraleetharan, Hellyer; performed data analysis: Wang, Muraleetharan, Hellyer; wrote or contributed to the writing of the manuscript: Wang, Hellyer, Gregory

## Conflict of interest statement

No author has an actual or perceived conflict of interest with the contents of this article

## Notes

### Competing Interest Statement

The authors have declared no competing interest.

## References

1. Klockgether T., Mariotti C., Paulson HL. Spinocerebellar ataxia. Nat Rev Dis Primer 2019;5(1):24. Doi: 10.1038/s41572-019-0074-3.

2. Sullivan R., Yau WY., O’Connor E., Houlden H. Spinocerebellar ataxia: an update. J Neurol 2019;266(2):533–44. Doi: 10.1007/s00415-018-9076-4.

3. Kasumu A., Bezprozvanny I. Deranged Calcium Signaling in Purkinje Cells and Pathogenesis in Spinocerebellar Ataxia 2 (SCA2) and Other Ataxias. The Cerebellum 2012;11(3):630–9. Doi: 10.1007/s12311-010-0182-9.

4. Robinson KJ., Watchon M., Laird AS. Aberrant Cerebellar Circuitry in the Spinocerebellar Ataxias. Front Neurosci 2020;14:707. Doi: 10.3389/fnins.2020.00707.

5. Paulson HL., Shakkottai VG., Clark HB., Orr HT. Polyglutamine spinocerebellar ataxias — from genes to potential treatments. Nat Rev Neurosci 2017;18(10):613–26. Doi: 10.1038/nrn.2017.92.

6. Gregory KJ., Goudet C. International Union of Basic and Clinical Pharmacology. CXI. Pharmacology, Signaling, and Physiology of Metabotropic Glutamate Receptors. Pharmacol Rev 2021;73(1):521–69. Doi: 10.1124/pr.119.019133.

7. Sugiyama H., Ito I., Hirono C. A new type of glutamate receptor linked to inositol phospholipid metabolism. Nature 1987;325(6104):531–3. Doi: 10.1038/325531a0.

8. Aiba A., Kano M., Chen C., Stanton ME., Fox GD., Herrup K., et al. Deficient cerebellar long-term depression and impaired motor learning in mGluR1 mutant mice. Cell 1994;79(2):377–88.

9. Conquet F., Bashir ZI., Davies CH., Daniel H., Ferraguti F., Bordi F., et al. Motor deficit and impairment of synaptic plasticity in mice lacking mGluR1. Nature 1994;372(6503):237–43. Doi: 10.1038/372237a0.

10. Kano M., Hashimoto K., Kurihara H., Watanabe M., Inoue Y., Aiba A., et al. Persistent multiple climbing fiber innervation of cerebellar Purkinje cells in mice lacking mGluR1. Neuron 1997;18(1):71–9. Doi: 10.1016/s0896-6273(01)80047-7.

11. Ichise T., Kano M., Hashimoto K., Yanagihara D., Nakao K., Shigemoto R., et al. mGluR1 in cerebellar Purkinje cells essential for long-term depression, synapse elimination, and motor coordination. Science 2000;288(5472):1832–5. Doi: 10.1126/science.288.5472.1832.

12. Guergueltcheva V., Azmanov DN., Angelicheva D., Smith KR., Chamova T., Florez L., et al. Autosomal-Recessive Congenital Cerebellar Ataxia Is Caused by Mutations in Metabotropic Glutamate Receptor 1. Am J Hum Genet 2012;91(3):553–64. Doi: 10.1016/j.ajhg.2012.07.019.

13. Davarniya B., Hu H., Kahrizi K., Musante L., Fattahi Z., Hosseini M., et al. The Role of a Novel TRMT1 Gene Mutation and Rare GRM1 Gene Defect in Intellectual Disability in Two Azeri Families. PLOS ONE 2015;10(8):e0129631. Doi: 10.1371/journal.pone.0129631.

14. Watson LM., Bamber E., Schnekenberg RP., Williams J., Bettencourt C., Lickiss J., et al. Dominant Mutations in GRM1 Cause Spinocerebellar Ataxia Type 44. Am J Hum Genet 2017;101(3):451–8. Doi: 10.1016/j.ajhg.2017.08.005.

15. Cabet S., Putoux A., Carneiro M., Labalme A., Sanlaville D., Guibaud L., et al. A novel truncating variant p.(Arg297*) in the GRM1 gene causing autosomal-recessive cerebellar ataxia with juvenile-onset. Eur J Med Genet 2019;62(10):103726. Doi: 10.1016/j.ejmg.2019.103726.

16. Ngo KJ., Rexach JE., Lee H., Petty LE., Perlman S., Valera JM., et al. A diagnostic ceiling for exome sequencing in cerebellar ataxia and related neurological disorders. Hum Mutat 2020;41(2):487–501. Doi: 10.1002/humu.23946.

17. Rasool IG., Zahoor MY., Iqbal M., Anjum AA., Ashraf F., Abbas HQ., et al. Whole exome sequencing revealed novel variants in consanguineous Pakistani families with intellectual disability. Genes Genomics 2021;43(5):503–12. Doi: 10.1007/s13258-021-01070-7.

18. Yousaf H., Fatima A., Ali Z., Baig SM., Toft M., Iqbal Z. A Novel Nonsense Variant in GRM1 Causes Autosomal Recessive Spinocerebellar Ataxia 13 in a Consanguineous Pakistani Family. Genes 2022;13(9):1667. Doi: 10.3390/genes13091667.

19. Protasova MS., Andreeva TV., Klyushnikov SA., Illarioshkin SN., Rogaev EI. Genetic Variant in GRM1 Underlies Congenital Cerebellar Ataxia with No Obvious Intellectual Disability. Int J Mol Sci 2023;24(2):1551. Doi: 10.3390/ijms24021551.

20. Serra HG., Byam CE., Lande JD., Tousey SK., Zoghbi HY., Orr HT. Gene profiling links SCA1 pathophysiology to glutamate signaling in Purkinje cells of transgenic mice. Hum Mol Genet 2004;13(20):2535–43. Doi: 10.1093/hmg/ddh268.

21. Notartomaso S., Zappulla C., Biagioni F., Cannella M., Bucci D., Mascio G., et al. Pharmacological enhancement of mGlu1 metabotropic glutamate receptors causes a prolonged symptomatic benefit in a mouse model of spinocerebellar ataxia type 1. Mol Brain 2013;6(1):48. Doi: 10.1186/1756-6606-6-48.

22. Armbrust KR., Wang X., Hathorn TJ., Cramer SW., Chen G., Zu T., et al. Mutant -III Spectrin Causes mGluR1 Mislocalization and Functional Deficits in a Mouse Model of Spinocerebellar Ataxia Type 5. J Neurosci 2014;34(30):9891–904. Doi: 10.1523/JNEUROSCI.0876-14.2014.

23. Konno A., Shuvaev AN., Miyake N., Miyake K., Iizuka A., Matsuura S., et al. Mutant Ataxin-3 with an Abnormally Expanded Polyglutamine Chain Disrupts Dendritic Development and Metabotropic Glutamate Receptor Signaling in Mouse Cerebellar Purkinje Cells. The Cerebellum 2014;13(1):29–41. Doi: 10.1007/s12311-013-0516-5.

24. Power EM., Morales A., Empson RM. Prolonged Type 1 Metabotropic Glutamate Receptor Dependent Synaptic Signaling Contributes to Spino-Cerebellar Ataxia Type 1. J Neurosci 2016;36(18):4910–6. Doi: 10.1523/JNEUROSCI.3953-15.2016.

25. Meera P., Pulst S., Otis T. A positive feedback loop linking enhanced mGluR function and basal calcium in spinocerebellar ataxia type 2. ELife 2017;18(6):e26377. Doi: 10.7554/eLife.26377.

26. Conn PJ., Lindsley CW., Meiler J., Niswender CM. Opportunities and challenges in the discovery of allosteric modulators of GPCRs for treating CNS disorders. Nat Rev Drug Discov 2014;13(9):692–708. Doi: 10.1038/nrd4308.

27. Ai N., Wood RD., Welsh WJ. Identification of Nitazoxanide as a Group I Metabotropic Glutamate Receptor Negative Modulator for the Treatment of Neuropathic Pain: An In Silico Drug Repositioning Study. Pharm Res 2015. Doi: 10.1007/s11095-015-1665-7.

28. Chen Y., Goudet C., Pin J-P., Conn PJ. N-{4-Chloro-2-[(1,3-dioxo-1,3-dihydro-2H-isoindol-2-yl)methyl]phenyl}-2-hydroxybenzamide (CPPHA) acts through a novel site as a positive allosteric modulator of group 1 metabotropic glutamate receptors. Mol Pharmacol 2008;73(3):909–18. Doi: 10.1124/mol.107.040097.

29. Hemstapat K., de Paulis T., Chen Y., Brady AE., Grover VK., Alagille D., et al. A novel class of positive allosteric modulators of metabotropic glutamate receptor subtype 1 interact with a site distinct from that of negative allosteric modulators. Mol Pharmacol 2006;70(2):616–26. Doi: 10.1124/mol.105.021857.

30. Knoflach F., Mutel V., Jolidon S., Kew JN., Malherbe P., Vieira E., et al. Positive allosteric modulators of metabotropic glutamate 1 receptor: characterization, mechanism of action, and binding site. Proc Natl Acad Sci U S A 2001;98(23):13402–7. Doi: 10.1073/pnas.231358298.

31. O’Brien JA., Lemaire W., Wittmann M., Jacobson MA., Ha SN., Wisnoski DD., et al. A novel selective allosteric modulator potentiates the activity of native metabotropic glutamate receptor subtype 5 in rat forebrain. J Pharmacol Exp Ther 2004;309(2):568–77. Doi: 10.1124/jpet.103.061747.

32. Vieira E., Huwyler J., Jolidon S., Knoflach F., Mutel V., Wichmann J. Fluorinated 9H-xanthene-9-carboxylic acid oxazol-2-yl-amides as potent, orally available mGlu1 receptor enhancers. Bioorg Med Chem Lett 2009;19(6):1666–9. Doi: 10.1016/j.bmcl.2009.01.108.

33. Lavreysen H., Wouters R., Bischoff F., Nóbrega Pereira S., Langlois X., Blokland S., et al. JNJ16259685, a highly potent, selective and systemically active mGlu1 receptor antagonist. Neuropharmacology 2004;47(7):961–72. Doi: 10.1016/j.neuropharm.2004.08.007.

34. Lavreysen H., Janssen C., Bischoff F., Langlois X., Leysen JE., Lesage ASJ. [3H]R214127: a novel high-affinity radioligand for the mGlu1 receptor reveals a common binding site shared by multiple allosteric antagonists. Mol Pharmacol 2003;63(5):1082–93. Doi: 10.1124/mol.63.5.1082.

35. Lovell KM., Felts AS., Rodriguez AL., Venable DF., Cho HP., Morrison RD., et al. N-Acyl-N’-arylpiperazines as negative allosteric modulators of mGlu1: identification of VU0469650, a potent and selective tool compound with CNS exposure in rats. Bioorg Med Chem Lett 2013;23(13):3713–8. Doi: 10.1016/j.bmcl.2013.05.020.

36. Brakeman PR., Lanahan AA., O’Brien R., Roche K., Barnes CA., Huganir RL., et al. Homer: a protein that selectively binds metabotropic glutamate receptors. Nature 1997;386(6622):284–8. Doi: 10.1038/386284a0.

37. Ciruela F., Robbins MJ., Willis AC., McIlhinney R a. J. Interactions of the C Terminus of Metabotropic Glutamate Receptor Type 1α with Rat Brain Proteins. J Neurochem 1999;72(1):346–54. Doi: 10.1046/j.1471-4159.1999.0720346.x.

38. Ciruela F., McIlhinney RA. Metabotropic glutamate receptor type 1alpha and tubulin assemble into dynamic interacting complexes. J Neurochem 2001;76(3):750–7. Doi: 10.1046/j.1471-4159.2001.00099.x.

39. Kitano J., Kimura K., Yamazaki Y., Soda T., Shigemoto R., Nakajima Y., et al. Tamalin, a PDZ domain-containing protein, links a protein complex formation of group 1 metabotropic glutamate receptors and the guanine nucleotide exchange factor cytohesins. J Neurosci Off J Soc Neurosci 2002;22(4):1280–9. Doi: 10.1523/JNEUROSCI.22-04-01280.2002.

40. Tu JC., Xiao B., Naisbitt S., Yuan JP., Petralia RS., Brakeman P., et al. Coupling of mGluR/Homer and PSD-95 complexes by the Shank family of postsynaptic density proteins. Neuron 1999;23(3):583–92. Doi: 10.1016/s0896-6273(00)80810-7.

41. Tu JC., Xiao B., Yuan JP., Lanahan AA., Leoffert K., Li M., et al. Homer Binds a Novel Proline-Rich Motif and Links Group 1 Metabotropic Glutamate Receptors with IP3 Receptors. Neuron 1998;21(4):717–26. Doi: 10.1016/S0896-6273(00)80589-9.

42. Uhlen M., Oksvold P., Fagerberg L., Lundberg E., Jonasson K., Forsberg M., et al. Towards a knowledge-based Human Protein Atlas. Nat Biotechnol 2010;28(12):1248–50. Doi: 10.1038/nbt1210-1248.

43. Vincent K., Cornea VM., Jong Y-JI., Laferrière A., Kumar N., Mickeviciute A., et al. Intracellular mGluR5 plays a critical role in neuropathic pain. Nat Commun 2016;7(1):10604. Doi: 10.1038/ncomms10604.

44. Ciruela F., Escriche M., Burgueno J., Angulo E., Casado V., Soloviev MM., et al. Metabotropic glutamate 1alpha and adenosine A1 receptors assemble into functionally interacting complexes. J Biol Chem 2001;276(21):18345–51. Doi: 10.1074/jbc.M006960200.

45. Lai TKY., Zhai D., Su P., Jiang A., Boychuk J., Liu F. The receptor-receptor interaction between mGluR1 receptor and NMDA receptor: a potential therapeutic target for protection against ischemic stroke. FASEB J 2019;33(12):14423–39. Doi: 10.1096/fj.201900417R.

46. Yang JH., Mao L-M., Choe ES., Wang JQ. Synaptic ERK2 Phosphorylates and Regulates Metabotropic Glutamate Receptor 1 In Vitro and in Neurons. Mol Neurobiol 2017;54(9):7156–70. Doi: 10.1007/s12035-016-0225-4.

47. White RR., Kwon Y-G., Taing M., Lawrence DS., Edelman AM. Definition of Optimal Substrate Recognition Motifs of Ca2+-Calmodulin-dependent Protein Kinases IV and II Reveals Shared and Distinctive Features *. J Biol Chem 1998;273(6):3166–72. Doi: 10.1074/jbc.273.6.3166.

48. Pippig S., Andexinger S., Daniel K., Puzicha M., Caron MG., Lefkowitz RJ., et al. Overexpression of beta-arrestin and beta-adrenergic receptor kinase augment desensitization of beta 2-adrenergic receptors. J Biol Chem 1993;268(5):3201–8. Doi: 10.1016/S0021-9258(18)53678-4.

49. Abreu N., Acosta-Ruiz A., Xiang G., Levitz J. Mechanisms of differential desensitization of metabotropic glutamate receptors. Cell Rep 2021;35(4):109050. Doi: 10.1016/j.celrep.2021.109050.

50. Desai MA., Burnett JP., Mayne NG., Schoepp DD. Pharmacological characterization of desensitization in a human mGlu1 alpha-expressing non-neuronal cell line co-transfected with a glutamate transporter. Br J Pharmacol 1996;118(6):1558–64. Doi: 10.1111/j.1476-5381.1996.tb15574.x.

51. Hellyer SD., Albold S., Sengmany K., Singh J., Leach K., Gregory KJ. Metabotropic glutamate receptor 5 (mGlu5)-positive allosteric modulators differentially induce or potentiate desensitization of mGlu5 signaling in recombinant cells and neurons. J Neurochem 2019;151(3):301–15. Doi: 10.1111/jnc.14844.

52. Iacovelli L., Salvatore L., Capobianco L., Picascia A., Barletta E., Storto M., et al. Role of G Protein-coupled Receptor Kinase 4 and ॆ-Arrestin 1 in Agonist-stimulated Metabotropic Glutamate Receptor 1 Internalization and Activation of Mitogen-activated Protein Kinases*. J Biol Chem 2003;278(14):12433–42. Doi: 10.1074/jbc.M203992200.

53. Ibrahim KS., Abd-Elrahman KS., Mestikawy SE., Ferguson SSG. Targeting Vesicular Glutamate Transporter Machinery: Implications on Metabotropic Glutamate Receptor 5 Signaling and Behavior. Mol Pharmacol 2020;98(4):314–27. Doi: 10.1124/molpharm.120.000089.

54. Doornbos MLJ., Van der Linden I., Vereyken L., Tresadern G., IJzerman AP., Lavreysen H., et al. Constitutive activity of the metabotropic glutamate receptor 2 explored with a whole-cell label-free biosensor. Biochem Pharmacol 2018;152:201–10. Doi: 10.1016/j.bcp.2018.03.026.

55. Hisatsune C., Hamada K., Mikoshiba K. Ca2+ signaling and spinocerebellar ataxia. Biochim Biophys Acta BBA - Mol Cell Res 2018;1865(11):1733–44. Doi: 10.1016/j.bbamcr.2018.05.009.

56. Prestori F., Moccia F., D’Angelo E. Disrupted Calcium Signaling in Animal Models of Human Spinocerebellar Ataxia (SCA). Int J Mol Sci 2019;21(1):216. Doi: 10.3390/ijms21010216.

57. Lei T., Hu Z., Ding R., Chen J., Li S., Zhang F., et al. Exploring the Activation Mechanism of a Metabotropic Glutamate Receptor Homodimer via Molecular Dynamics Simulation. ACS Chem Neurosci 2020;11(2):133–45. Doi: 10.1021/acschemneuro.9b00425.

58. Koehl A., Hu H., Feng D., Sun B., Zhang Y., Robertson MJ., et al. Structural insights into the activation of metabotropic glutamate receptors. Nature 2019;566(7742):79–84. Doi: 10.1038/s41586-019-0881-4.

59. Zhang J., Qu L., Wu L., Tang X., Luo F., Xu W., et al. Structural insights into the activation initiation of full-length mGlu1. Protein Cell 2021;12(8):662–7. Doi: 10.1007/s13238-020-00808-5.

60. Ashiru O., Howe JD., Butters TD. Nitazoxanide, an antiviral thiazolide, depletes ATP-sensitive intracellular Ca2+ stores. Virology 2014;462–463:135–48. Doi: 10.1016/j.virol.2014.05.015.

61. Muraleetharan A., Wang Y., Rowe MC., Gould A., Gregory KJ., Hellyer SD. Rigorous Characterization of Allosteric Modulation of the Human Metabotropic Glutamate Receptor 1 Reveals Probe- and Assay-Dependent Pharmacology. Mol Pharmacol 2023;103(6):325–38. Doi: 10.1124/molpharm.122.000664.

62. Sheffler DJ., Conn PJ. Allosteric potentiators of metabotropic glutamate receptor subtype 1a differentially modulate independent signaling pathways in baby hamster kidney cells. Neuropharmacology 2008;55(4):419–27. Doi: 10.1016/j.neuropharm.2008.06.047.

63. Emery AC., DiRaddo JO., Miller E., Hathaway HA., Pshenichkin S., Takoudjou GR., et al. Ligand bias at metabotropic glutamate 1a receptors: molecular determinants that distinguish β-arrestin-mediated from G protein-mediated signaling. Mol Pharmacol 2012;82(2):291–301. Doi: 10.1124/mol.112.078444.

64. Hathaway HA., Pshenichkin S., Grajkowska E., Gelb T., Emery AC., Wolfe BB., et al. Pharmacological characterization of mGlu1 receptors in cerebellar granule cells reveals biased agonism. Neuropharmacology 2015;93:199–208. Doi: 10.1016/j.neuropharm.2015.02.007.

