## Supplementary Tables and Figures for "SCA44- and SCAR13-associated *GRM1* mutations affect metabotropic glutamate receptor 1 function through distinct mechanisms"

**Table S1. Mutagenesis primers used in the current study**

| Primer | Direction | Sequence (5’ to 3’) |
| --- | --- | --- |
| Insertion of cMyc tag | Forward | gagcaaaagctcatttctgaagaggacttgctcgaaaaccctaatt |
|  | Reverse | caagtcctcttcagaaatgagcttttgctccaggtgcccgggcaac |
| c1360t_g1362c (L454F) | Forward | cactcctatgaaggaggatttaatgaagaaatccagaagtttagaacc |
|  | Reverse | ggttctaaacttctggatttcttcattaaatcctccttcataggagtg |
| del2652-54 (N885del) | Forward | gaaagctggagccggcgccaattctaacgg |
|  | Reverse | ccgttagaattggcgccggctccagctttc |
| 3165dup (G1056Rfs*49) | Forward | ccaggtcccccttggcccggcgagaactg |
|  | Reverse | cagttctcgccgggccaagggggacctgg |
| a2375g (Y792C) | Forward | ctccggcattggagcaaattttatcggagtgtgcaatac |
|  | Reverse | gtattgcacactccgataaaatttgctccaatgccggag |
| a785g (Y262C) | Forward | gccagataatgcaggtagtacacatggtaaaagcgatgtac |
|  | Reverse | gtacatcgcttttaccatgtgtactacctgcattatctggc |

**Table S2. Receptor expression of WT and mutant mGlu_1_ in Flp-In TRex HEK293 cells induced with different concentration of tetracycline.** Data are mean ± S.E.M. pooled from the indicated number of independent experiments.

|  | Tetracycline concentration | | | |
| --- | --- | --- | --- | --- |
|  | 0ng/ml | 30ng/ml | 1μg/ml | n |
| WT | 1.40 ± 0.31 | 18.56 ± 7.96 | 33.70 ± 12.98 | 6 |
| Y262C | 1.00 ± 0.19 | 5.97 ± 3.20 | 8.94 ± 4.37 | 5 |
| L454F | 1.58 ± 0.80 | 6.15 ± 2.89 | 14.03 ± 5.92 | 5 |
| Y792C | 1.93 ± 0.93 | 12.27 ± 8.01 | 27.18 ± 14.64 | 5 |
| N885del | 0.99 ± 0.16 | 5.58 ± 0.92 | 15.94 ± 3.39 | 4 |
| G1056Rfs*49 | 1.15 ± 0.45 | 1.86 ± 0.91 | 3.90 ± 2.94 | 6 |

NB: surface expression is normalised as fold-over fluorescence of non-transfected Flp-In TREx HEK293 cells

**Table S3.** Orthosteric ligand potency (pEC_50_) and maximal response (E_max_) estimates for iCa^2+^ mobilisation and IP_1_ accumulation in Flp-In TREx HEK293 cells expressing wild type and mutant mGlu_1_ at expression levels induced by 30ng/ml of tetracycline. Data are mean ± S.E.M of indicated number (n) of independent experiments performed in duplicate.

|  | iCa^2+^ mobilisation | | | | | | IP_1_ accumulation | | | | | |
| --- | --- | --- | --- | --- | --- | --- | --- | --- | --- | --- | --- | --- |
|  | glutamate | | | quisqualate | | | quisqualate | | | DHPG | | |
| Mutants | pEC_50_^a^ | E_max_^b^ | n | pEC_50_^a^ | E_max_^b^ | n | pEC_50_^a^ | E_max_^c^ | n | pEC_50_^a^ | E_max_^c^ | n |
| WT | 6.22 ± 0.09^e^ | 50.6 ± 2.8 | 13 | 7.42 ± 0.09 | 47.0 ± 3.6 | 12 | 6.83 ± 0.12 | 2.78 ± 0.21 | 11 | 5.55 ± 0.08 | 2.56 ± 0.47 | 6 |
| Y262C | 5.97 ± 0.12^e^ | 43.7 ± 3.6 | 11 | 7.22 ± 0.12 | 38.5 ± 5.7 | 12 | 6.83 ± 0.17 | 2.75 ± 0.24 | 17 | n.d. | 2.18 ± 0.15 | 5 |
| L454F | 5.51 ± 0.12^d,e^ | 61.6 ± 6.3 | 17 | 7.01 ± 0.11 | 47.5 ± 4.0 | 11 | 6.45 ± 0.09 | 3.01 ± 0.29 | 16 | n.d. | 2.14 ± 0.25 | 8 |
| Y792C | 6.34 ± 0.10^e^ | 45.5 ± 4.9 | 16 | 7.68 ± 0.11 | 42.7 ± 6.0 | 10 | 6.56 ± 0.11^e^ | 1.60 ± 0.08^d^ | 14 | n.d. | 1.57 ± 0.17^d^ | 6 |
| N885del | 5.49 ± 0.10^d,e^ | 51.2 ± 6.7^e^ | 16 | 6.80 ± 0.17^d^ | 25.5 ± 4.8^d^ | 13 | 6.91 ± 0.07 | 2.59 ± 0.29 | 15 | n.d. | 2.27 ± 0.20 | 7 |
| G1056Rfs*49 | n.r. | n.r. | 5 | n.r. | n.r. | 4 | 6.32 ± 0.17 | 1.52 ± 0.22^d^ | 6 | n.r | n.r. | 4 |

n.r. no response, n.d. not determined due to poor curve fit

^a^ The negative logarithm of the molar concentration of agonist required to yield a half maximal response.

^b^ E_max_ of iCa^2+^ mobilisation is expressed as percentage of ionomycin response

^c^ E_max_ of IP_1_ accumulation is expressed as fold of basal. Due to a lack of plateau for DHPG curves, this represents the response at 30µM DHPG for all mutant receptors

^d^ Significantly different from mGlu_1_ WT response (p < 0.05, one-way ANOVA, Šídák's post-test).

^e^ significantly different from iCa^2+^ response to quisqualate (p < 0.05, one-way ANOVA, Šídák's post-test).

**Table S4.** Orthosteric ligand functional affinity (pK_A_) and efficacy (log_τ_) estimates derived from receptor titration iCa^2+^ mobilisation experiments in Flp-In TREx HEK293 cells expressing different levels of wild type and mutant mGlu_1_. Data are mean ± S.E.M of indicated number (n) of independent experiments performed in duplicate.

|  | glutamate | | | | quisqualate | | | |
| --- | --- | --- | --- | --- | --- | --- | --- | --- |
| Mutants | pK_A_^a^ | Log_τ_ basal^b^ | Log_τ_ 30ng^c^ | n | pK_A_^a^ | Log_τ_ basal | Log_τ_ 30ng | n |
| WT | 5.39 ± 0.08 | -0.21 ± 0.07 | 0.64 ± 0.07^e^ | 5 | 7.08 ± 0.11^f^ | -0.63 ± 0.11 | 0.12 ± 0.05^e,f^ | 6 |
| Y262C | 5.38 ± 0.06 | -0.17 ± 0.05 | 0.45 ± 0.05^e^ | 5 | 6.76 ± 0.09^f^ | -0.68 ± 0.12^f^ | 0.38 ± 0.05^e^ | 6 |
| L454F | 4.91 ± 0.15^d^ | n.d. | 0.34 ± 0.11 | 5 | 6.38 ± 0.08^d,f^ | n.d. | -0.06 ± 0.04^e^ | 5 |
| Y792C | 5.84 ± 0.10 | -0.50 ± 0.11 | 0.37 ± 0.07^e^ | 6 | 7.16 ± 0.17^f^ | -0.22 ± 0.13 | 0.47 ± 0.10^e^ | 5 |
| N885del | 5.09 ± 0.12 | -0.88 ± 0.18^d^ | 0.07 ± 0.07^d,e^ | 7 | 6.56 ± 0.11^d,f^ | -0.87 ± 0.10 | -0.32 ± 0.05^d,e^ | 7 |
| G1056Rfs*49 | n.r. | n.r. | n.r. | 5 | n.r. | n.r. | n.r. | 5 |

^a^ The negative logarithm of the equilibrium dissociation constant of orthosteric agonists. Curves were fitted globally to n= 4-7 independent experiments

^b^ intrinsic efficacy of the orthosteric agonist under basal receptor expression conditions (i.e. no tetracycline)

^c^ intrinsic efficacy of the orthosteric agonist under comparable maximum receptor expression conditions (i.e. 30ng/ml tetracycline)

^d^ Significantly different from mGlu_1_ WT response (p < 0.05, one-way ANOVA, Šídák's post-test).

^e^ Significantly different from basal response for matched agonist (p < 0.05, one-way ANOVA, Šídák's post-test).

^f^ Significantly different from glutamate estimate (p < 0.05, one-way ANOVA, Šídák's post-test).

**Table S5.** Potency, maximum response, affinity and cooperativity estimates for PAM modulation of orthosteric agonist induced iCa^2+^ mobilisation and IP_1_ accumulation in Flp-In TRex HEK293 expressing mGlu_1_ WT and loss-of-function mutations L454F and N885del. Data are mean ± SEM of indicated number (n) of independent experiments performed in duplicate.

|  |  | iCa^2+^ mobilisation | | | | | | | | | | IP_1_ accumulation | | | | | | | |
| --- | --- | --- | --- | --- | --- | --- | --- | --- | --- | --- | --- | --- | --- | --- | --- | --- | --- | --- | --- |
|  |  | + glutamate EC_20_ | | | | | + quisqualate EC_20_ | | | | | PAM alone | | | + quisqualate EC_20_ | | | | |
|  |  | pMod_50_^a^ | Mod_max_^b^ | pK_b_^c^ | Log β^d^ | n | pMod_50_ | Mod_max_ | pK_b_ | Log β | n | pEC_50_ | E_max_^e^ | n | pMod_50_ | Mod_max_^e^ | pK_b_ | Log β | n |
| WT | CPPHA | n.d. | 15.1 ±2.8 | n.d | n.d | 7 | n.d. | 15.4 ±4.3 | n.d | n.d | 7 | n.d. | 0.50 ±0.08 | 7 | n.d | 0.52 ±0.12 | n.d | n.d | 5 |
|  | Ro 67-4853 | 7.26 ±0.27 | 31.5 ±4.6 | 6.92 ±0.39 | 0.78 ±0.17 | 7 | 7.46 ±0.10 | 27.7 ±5.0 | 7.02 ±0.47 | 0.35 ±0.13 | 7 | n.d | 0.89 ±0.21 | 7 | 6.25 ±0.20^f^ | 0.89 ±0.07 | 5.61 ±0.22^f^ | 0^i^ | 5 |
| L454F | CPPHA | n.d. | 16.1 ±6.7 | n.d | n.d | 8 | n.d. | 12.3 ±2.1 | n.d | n.d | 6 | n.r. | n.r. | 5 | n.d. | 0.27 ±0.10 | n.d | n.d | 8 |
|  | Ro 67-4853 | n.d. | 30.0 ±9.2 | n.d | n.d | 7 | 7.70 ±0.30 | 15.8 ±6.9 | 7.00 ±0.54 | 0.56 ±0.16 | 5 | n.d. | 0.17 ±0.04^h^ | 5 | 5.83 ±0.11^f^ | 0.90 ±0.13^g^ | 6.05  ±0.28^f^ | 0.45 ±0.07 | 8 |
| N885 del | CPPHA | n.d. | 16.8 ±7.6 | n.d | n.d | 6 | n.d. | 8.7  ±4.5 | n.d | n.d | 6 | n.d. | 0.37 ±0.04 | 6 | n.d. | 0.78 ±0.22 | n.d | n.d | 6 |
|  | Ro 67-4853 | 6.64 ±0.37 | 18.1 ±6.5 | 7.17 ±0.54 | 0.56 ±0.18 | 5 | n.d. | 6.2  ±2.0 | n.d | n.d | 6 | n.d. | 0.78 ±0.15 | 6 | 6.21 ±0.15^f^ | 1.06 ±0.33 | n.d. | n.d | 6 |

n.d. not determined due to poor curve fit; n.r. no response

^a^ The negative logarithm of the molar concentration of agonist required to yield a half maximal response in the presence of EC_20_ agonist (pMod_50_).

^b^ Mod_max_ of iCa^2+^ mobilisation is the span between EC_20_ baseline and the highest concentration of PAM (3µM for Ro 67-4853 and 10µM for CPPHA), expressed as percentage of ionomycin response. EC_20_ concentrations used are derived from concentration-response curves for orthosteric ligands.

^c^ pK_b_, negative logarithm of the equilibrium dissociation constant, derived from global fitting to operational model of allosterism

^d^ logβ, logarithm of the efficacy modulation factor, derived from global fitting to operational model of allosterism

^e^ E_max_/Mod_max_ of IP_1_ accumulation is the span between baseline (E_max_) or EC_20_ baseline (Mod_max_) and responses at highest concentration of PAM and are expressed as fold of basal. EC_20_ concentrations used are derived from concentration-response curves for orthosteric ligands

^f^ significantly different from modulation of quisqualate mediated iCa^2+^ mobilisation (p < 0.05, one-way ANOVA, Šídák's post-test)

^g^ significantly different from PAM alone in IP_1_ modulation (p < 0.05, one-way ANOVA, Šídák's post-test)

^h^ significantly different from WT (p < 0.05, one-way ANOVA, Šídák's post-test)

^i^ curves were best fit to a model in which cooperativity was neutral (i.e. logβ = 0), based on an extra sum-of-squares F-test

**Table S6.** Potency, maximum response, affinity and cooperativity estimates for NAM modulation of orthosteric agonist induced iCa^2+^ mobilisation and IP_1_ accumulation in Flp-In TRex HEK293 expressing mGlu_1_ WT and gain-of-function mutations Y262C and Y792C. Data are mean ± SEM of indicated number (n) of independent experiments performed in duplicate.

|  |  | iCa^2+^ mobilisation | | | | | | | | | | IP_1_ accumulation | | | | | | | |
| --- | --- | --- | --- | --- | --- | --- | --- | --- | --- | --- | --- | --- | --- | --- | --- | --- | --- | --- | --- |
|  |  | + glutamate EC_20_ | | | | | + quisqualate EC_20_ | | | | | modulator alone | | | + quisqualate EC_20_ | | | | |
|  |  | pMod_50_^a^ | Mod_max_^b^ | pK_b_^c^ | Log β^d^ | n | pMod_50_ | Mod_max_ | pK_b_ | Log β | n | pEC_50_ /  pIC_50_ | I_max_^e^ | n | pMod_50_ | Mod_max_^f^ | pK_b_ | Log β | n |
| WT | VU 0469650 | 7.80 ±0.19 | -37.5 ±6.2 | 8.40 ±0.17 | Full NAM | 7 | 7.67 ±0.11 | -33.9 ±4.3 | 7.86 ±0.17 | Full NAM | 7 | 8.44 ±0.35 | -0.15 ±0.01 | 5 | 8.17 ±0.51 | -57.1 ±7.0 | 8.39 ±0.37 | -0.41 ±0.11 | 5 |
|  | JNJ 16259685 | 8.78 ±0.17 | -40.4 ±6.6 | 9.49 ±0.16 | Full NAM | 7 | 8.65 ±0.13 | -36.2 ±4.8 | 8.91 ±0.17 | Full NAM | 7 | 8.65 ±0.26 | -0.15 ±0.01 | 5 | 7.73 ±0.36 | -103 ±26.0 | 7.88 ±0.20 | Full NAM | 5 |
| Y262C | VU 0469650 | 7.46 ±0.17 | -26.5 ±4.6 | 7.65 ±0.28 | Full NAM | 6 | 7.38 ±0.16 | -32.1 ±9.6 | 8.10 ±0.26 | Full NAM | 6 | 9.85 ±0.30^g^ | -0.21 ±0.03 | 5 | 8.30 ±0.40^i^ | -51.0 ±4.8 | 8.68 ±0.42 | -0.61 ±0.15 | 5 |
|  | JNJ 16259685 | 8.53 ±0.29 | -30.1 ±6.6 | 8.66 ±0.29 | Full NAM | 6 | 8.49 ±0.34 | -37.8 ±10.8 | 8.93 ±0.26 | Full NAM | 6 | 9.47 ±0.26 | -0.14 ±0.03 | 5 | 8.11 ±0.35^i^ | -81.5 ±7.3 | 8.31 ±0.26 | Full NAM | 5 |
|  | CPPHA | n.t. | n.t. | n.t. | n.t. |  | n.t. | n.t. | n.t. | n.t. |  | n.r. | n.r. | 5 | n.t. | n.t. | n.t. | n.t. |  |
|  | Ro 67-4853 | n.t. | n.t. | n.t. | n.t. |  | n.t. | n.t. | n.t. | n.t. |  | 6.46 ±0.13 | 0.34 ±0.03 | 5 | n.t. | n.t. | n.t. | n.t. |  |
| Y792C | VU 0469650 | 7.04 ±0.13 | -35.8 ±8.5 | 7.24 ±0.35^g^ | Full NAM | 7 | 7.30 ±0.12 | -36.6 ±5.1 | 7.94 ±0.18 | Full NAM | 5 | 9.09 ±0.07 | -0.42 ±0.03^g^ | 5 | 8.69 ±0.22^h^ | -73.7 ±18.5 | 8.51 ±0.24 | Full NAM | 5 |
|  | JNJ 16259685 | 8.38 ±0.24 | -40.8 ±11.0 | 8.54 ±0.33 | Full NAM | 7 | 8.58 ±0.38 | -40.4 ±6.4 | 9.05 ±0.18 | Full NAM | 5 | 9.02 ±0.13 | -0.46 ±0.04^g^ | 5 | 9.02 ±0.45^g^ | -98.3 ±25.7 | 9.53 ±0.28^g^ | Full NAM | 5 |
|  | CPPHA | n.t. | n.t. | n.t. | n.t. |  | n.t. | n.t. | n.t. | n.t. |  | n.d. | 0.30 ±0.09 | 6 | n.t. | n.t. | n.t. | n.t. |  |
|  | Ro 67-4853 | n.t. | n.t. | n.t. | n.t. |  | n.t. | n.t. | n.t. | n.t. |  | 6.52 ±0.15 | 0.57 ±0.08 | 6 | n.t. | n.t. | n.t. | n.t. |  |

n.r. no response, n.d. not determined due to poor curve fit

^a^ The negative logarithm of the molar concentration of agonist required to yield a half inhibitory response in the presence of EC_80_ agonist

^b^ Mod_max_ of iCa^2+^ mobilisation is the span between EC_80_ baseline and the bottom plateau of NAM curves, expressed as percentage of ionomycin response. EC_80_ concentrations used are derived from concentration-response curves for orthosteric ligands.

^c^ pK_b_, negative logarithm of the equilibrium dissociation constant, derived from global fitting to operational model of allosterism

^d^ logβ, logarithm of the efficacy modulation factor, derived from global fitting to operational model of allosterism

^e^ I_max_ of IP_1_ accumulation is the span between baseline and responses at highest concentration of modulator and are expressed as fold of basal.

^f^ Due to difference in signal window for Y792C compared to WT and Y262C, Mod_max_ of IP_1_ accumulation is presented the span between EC_80_ baseline and the bottom plateau of NAM curves in the presence of EC_80_, expressed as percentage of quisqualate response. EC_80_ concentrations used are derived from concentration-response curves for orthosteric ligands

^g^ significantly different from mGlu_1_ WT response (p < 0.05, one-way ANOVA, Šídák's post-test)

^h^ significantly different from modulation of quisqualate mediated iCa^2+^ mobilisation (p < 0.05, one-way ANOVA, Šídák's post-test)

^i^ significantly different from PAM alone in IP_1_ modulation (p < 0.05, one-way ANOVA, Šídák's post-test)

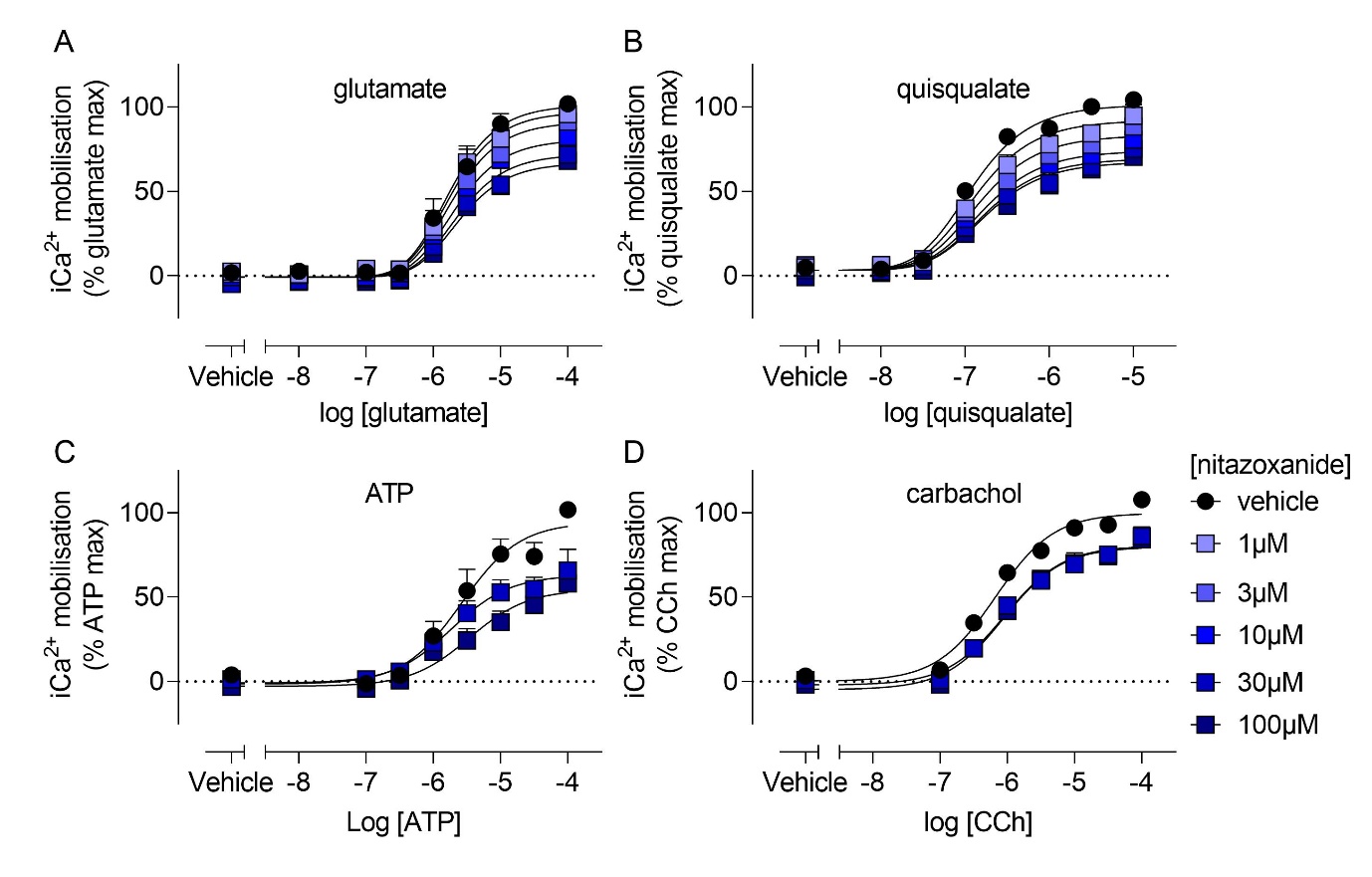

**Figure S1. Nitazoxanide has non-selective effects on cellular iCa^2+^ signalling.** Nitazoxanide reduces iCa^2+^ mobilisation induced by glutamate (A) and quisqualate (B) in HEK293A cells stably expressing mGlu_1_, and by ATP (C) and carbachol (D) in non-transfected HEK293A cells. Data represent mean + SEM of 5 experiments performed in duplicate. Error bars not shown lie within the dimensions of the symbol.

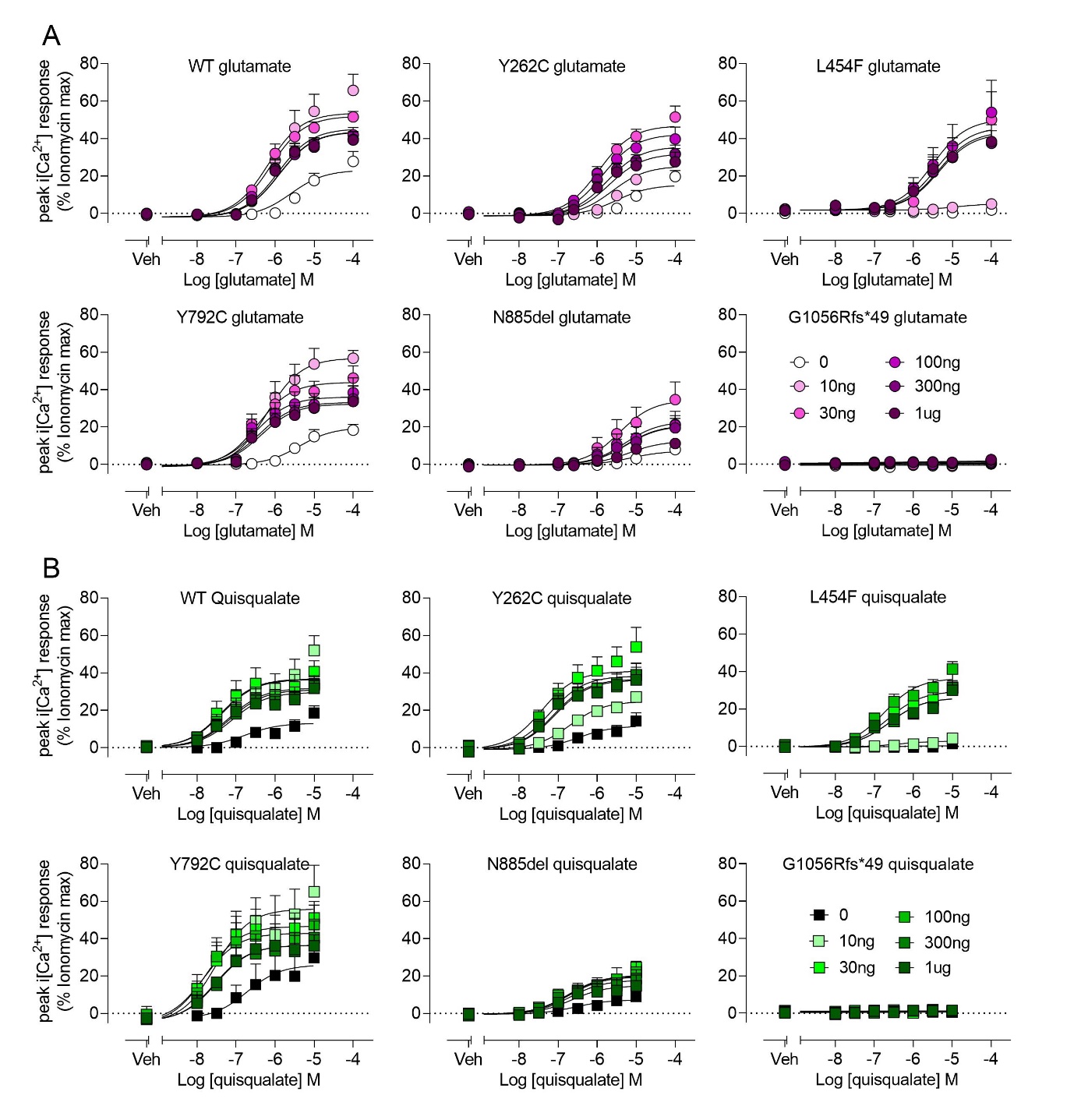

**Figure S2. Glutamate and quisqualate mediated iCa^2+^ mobilisation are differentially altered in mGlu_1_ mutants at varying concentration of tetracycline in an inducible Flp-In T-REx HEK293 cell line.** Expression of WT and mutant mGlu_1_ was induced by incubating Flp-In T-REx HEK293 cells with different concentrations of tetracycline overnight. Data are mean + S.E.M. from 5-6 experiments performed in duplicate. With the exception of L454F, all cell lines exhibit orthosteric agonist induced iCa^2+^ mobilisation in the absence of tetracycline. G0156Rfs*49 exhibits no response at any tetracycline concentration.
